# Atomistic Mechanism of Non-Canonical Voltage Gating in TREK K_2P_ Channels

**DOI:** 10.1101/2025.01.08.631886

**Authors:** Yessenbek K. Aldakul, Marcus Schewe, Carlos C. Diez, Songhwan Hwang, Thomas Baukrowitz, Han Sun

## Abstract

Two-pore-domain K^+^ (K_2P_) channels are essential regulators of cellular excitability and respond to various external stimuli that also include membrane potential for most K_2P_ channels. Voltage gating is particularly pronounced in members of the TREK/TRAAK subfamily. This voltage sensitivity was surprising, as K_2P_ channels lack a classical voltage sensing domain. Prior studies have attributed this non-canonical voltage sensing mechanism to the unique ion-flux gating properties of the K_2P_ selectivity filter (SF), where inward currents induce fast inactivation, while depolarization activates the SF. Here, we performed large-scale molecular dynamics (MD) simulations across various voltages to gain an atomistic understanding of ion-flux gating in TREK channels. Our analysis revealed an asymmetric stability difference in the SF, enabling water influx into the filter due to conformational flexibility on the extracellular side. Inward flux inactivation occurs when water entry halts ion permeation, followed by the unbinding of three K^+^ ions, consistent with gating charge analysis. Additionally, MD simulations of TREK-2 mutants and the TWIK-1 channel, which exhibited increased SF flexibility, showed augmented SF water occupancy, aligning with their electrophysiological phenotypes. Key experimental evidence of this mechanism was provided by electrophysiology measurements, which showed that high extracellular sucrose slowed ion-flux inactivation by reducing water entry into the SF. These findings uncover the atomistic mechanism of voltage gating in TREK K_2P_ channels and pave the way for exploring non-canonical voltage gating mechanisms in other ion channels.

## Introduction

Two-pore-domain K^+^ (K_2P_) channels control the resting potential and, thereby, membrane excitability in various cell types and organs. They are critically involved in a wide range of physiological functions, such as oxygen sensing, temperature sensitivity, pain perception, and play an important role in anesthesia and cardiac function (*1*). Structurally, different from other K^+^ selective channels, K_2P_ channels are formed as dimers with two pore-forming domains in each subunit. Among the 15 members of the K_2P_ channel family, atomistic structures have been resolved for TWIK-1 (*2*, *3*), TREK-1 (*4–6*), TREK-2 (*7*), TRAAK (*8–12*), TASK-1 (*13*, *14*), TASK-2 (*15*), TASK-3 (*14*, *16*), and THIK-1 channels (*17*, *18*). Previous work by us and others have revealed that the selectivity filter (SF) acts as the main gate in K_2P_ channels (*19–24*), with structural evidence of SF gating in TREK channels observed at low K^+^ concentration (*5*) and in TWIK and TASK channels at different pH levels (*3*, *16*). However, recent studies have identified an additional lower gate in TASK-1, referred to as the “X-gate” (*13*, *14*), and in THIK-1 termed as the “Y-gate” (*17*, *18*). Both gates create a constriction site within the ion permeation pathway in the cavity below the SF.

Although initially identified as background K^+^ channels, K_2P_ channels were later found to be subject to complex regulatory mechanisms, including membrane potential, mechanical forces, temperature alterations, (de-)phosphorylation, and membrane lipids like phosphatidylinositol-4,5-bisphosphate (PIP_2_) (*25–29*). Different from various stimuli that primarily act on the intracellular C-terminus and central cavity, our previous study suggested that voltage sensitivity arises from the SF (*19*). This finding indicates that the voltage gating mechanism in K_2P_ channels differs markedly from that in other voltage-gated cation channels, such as K_v_, Na_v_, Ca_v_, hyperpolarization-activated cyclic nucleotide-gated (HCN), and transient receptor potential (TRP) channels, where the voltage-sensing domain (VSD) controls pore opening in response to changes in membrane potential (Fig. 1A) (*30*, *31*). By contrast, in K_2P_ channels, voltage sensitivity is thought to emerge from an ion-flux-dependent gating process within the SF. The activation of the filter relies on the energy derived from the movement of ions into the SF and is thus powered by the electrochemical gradient. Central to this concept is the observation that inward ion-flux induces an inactivation process in the SF of most K_2P_ channels, most prominently in the TREK/TRAAK subfamily. Conversely, membrane potentials positive to the reversal potential drive ions into the inactivated SF, resulting in channel activation (Fig. 1B). However, the atomistic mechanism underlying voltage gating—in particular the voltage-dependent inactivation of the SF in K_2P_ channels— remains largely uncharted.

**Figure 1.**
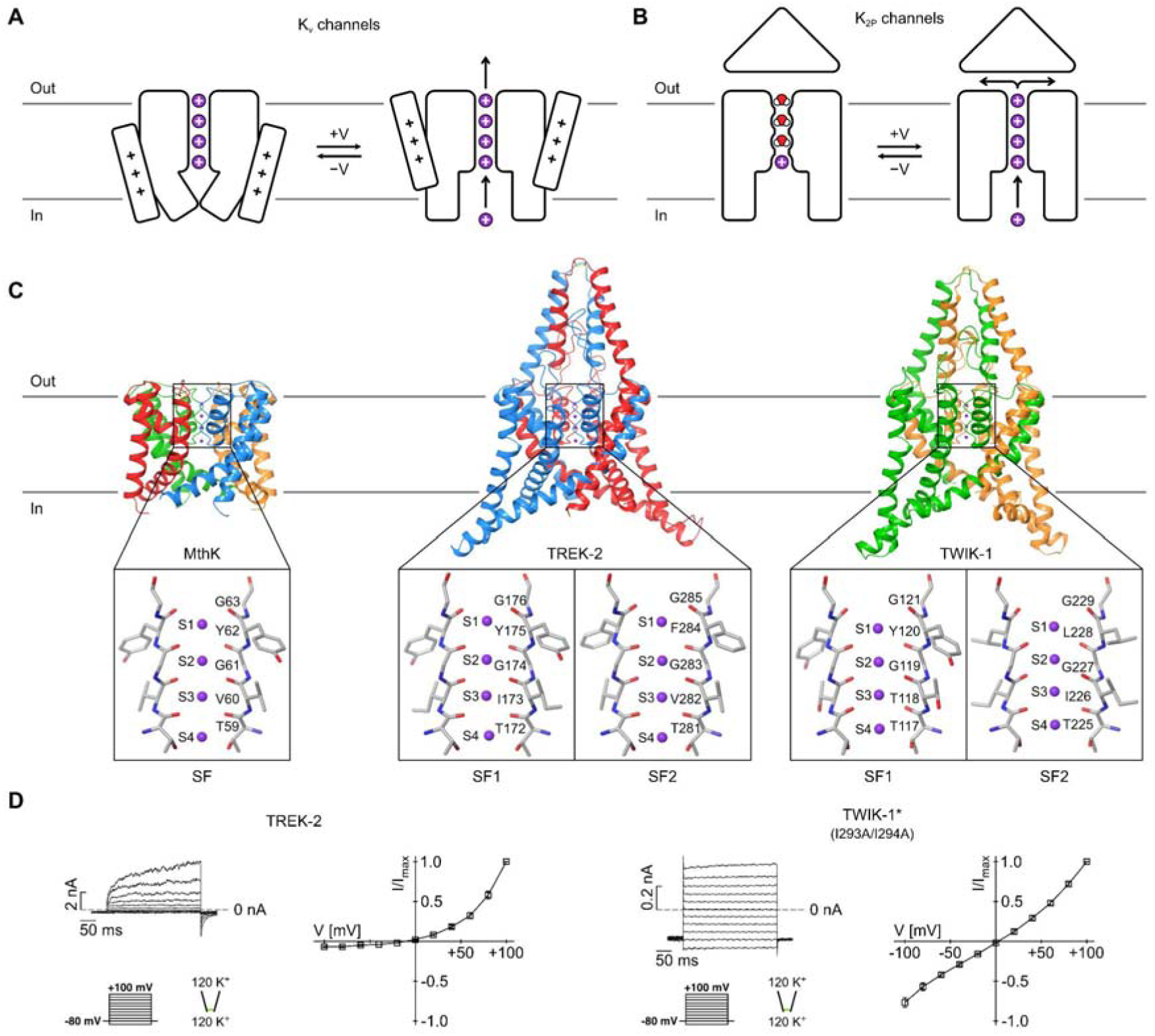
Voltage gating in K^+^ channels. (**A** and **B**) Schematic illustration of the canonical (A) and non-canonical (B) voltage gating mechanisms in K_v_ and K_2P_ channels, respectively. (**C**) Structures of MthK (PDB: 3LDC), TREK-2 (PDB: 4BW5), and TWIK-1 (PDB: 7SK0) with enlarged view of opposite SF subunits. The subunits of the channels are displayed in different colors and K^+^ ions are shown as purple spheres. (**D**) I-V curves of TREK-2 and TWIK-1* analyzed from current-voltage responses measured with the indicated voltage protocol in inside-out patches of oocytes in symmetrical high K^+^ [120 mM ex./120 mM int.] at pH 7.4. Analyzed electrophysiological data shown as mean ± SEM. The representative experiments were repeated (n > 6) with similar results.

To address this gap, we conducted large-scale atomistic molecular dynamics (MD) simulations totaling over 150 μs on three different types of K^+^ channels (Fig. 1C). Our primary focus was the voltage-gated TREK-2 K_2P_ channel (Fig. 1D), which shows strong ion-flux gating properties. For comparison, we also simulated the voltage-insensitive TWIK-1 K_2P_ channel, characterized by its unique low intrinsic activity among K_2P_ channels (Fig. 1D), with SF instability proposed as the main underlying cause (*32*). Additionally, we included MthK, a weakly inwardly-rectifying K^+^ channel (*33*), as another reference in our simulations. MthK exhibits a relatively high single-channel conductance and a canonical signature SF sequence.

We employed the MD-based Computational Electrophysiology (CompEL) approach (*34*) to simulate ion permeations under both positive and negative voltages for TREK-2, TWIK-1, MthK, as well as TREK-2 mutants. We demonstrate that the conformational instability of the extracellular side of the SF in TREK-2 initiates filter inactivation under negative voltages by enabling water influx, which was not observed in the simulations under positive voltages. MD simulations further revealed a fully inactive SF conformation with one K^+^ bound at the S4 K^+^ coordination site, validated by comparison of experimentally and computationally derived gating charges. We further show that SF inactivation associated with voltage gating results in additional conformational changes in the “TM3 glutamate network”, consistent with previous structural studies of TREK-1 at low K^+^ concentrations (*5*). Finally, the critical role of extracellular water in channel inactivation under negative voltages was confirmed through electrophysiological measurements, where the availability of extracellular water molecules was reduced by introducing sucrose.

## Results

### MD simulations of TREK-2, TWIK-1 and MthK under positive and negative voltages revealed distinct water occupancies in the SF

We performed atomistic MD simulations of TREK-2, TWIK-1, and the pore domain of MthK at 303 K under both positive and negative voltages (∼+200 mV or –200 mV, respectively, Table S1) within a single simulation box, following the CompEL protocol (*34*). The simulations were performed with the CHARMM36m force field (*35*). Under positive voltages, TREK-2 showed continuous K^+^ permeations (Fig. 2D, F, Fig. S2, Mov. 1, 2). While traversing the SF, K^+^ ions primarily occupied the S1-S4 sites, with notably smaller K^+^ occupancy at S3 (Fig. 2E). In contrast, under negative voltages, inward K^+^ permeations were observed during the initial period of the simulations. In most simulation runs, however, permeations halted within the first 1 μs (Fig. 2D, F, Fig. S2, Mov. 3, 4). This event was consistently accompanied by water influx, particularly occupying the S1 and S3 sites. As shown in Fig. 2E, water occupancy at S3 is significantly increased under negative voltages compared to positive voltages. Simultaneously, K^+^ occupancy at the S1 and S3 sites almost disappeared under negative voltages.

**Figure 2.**
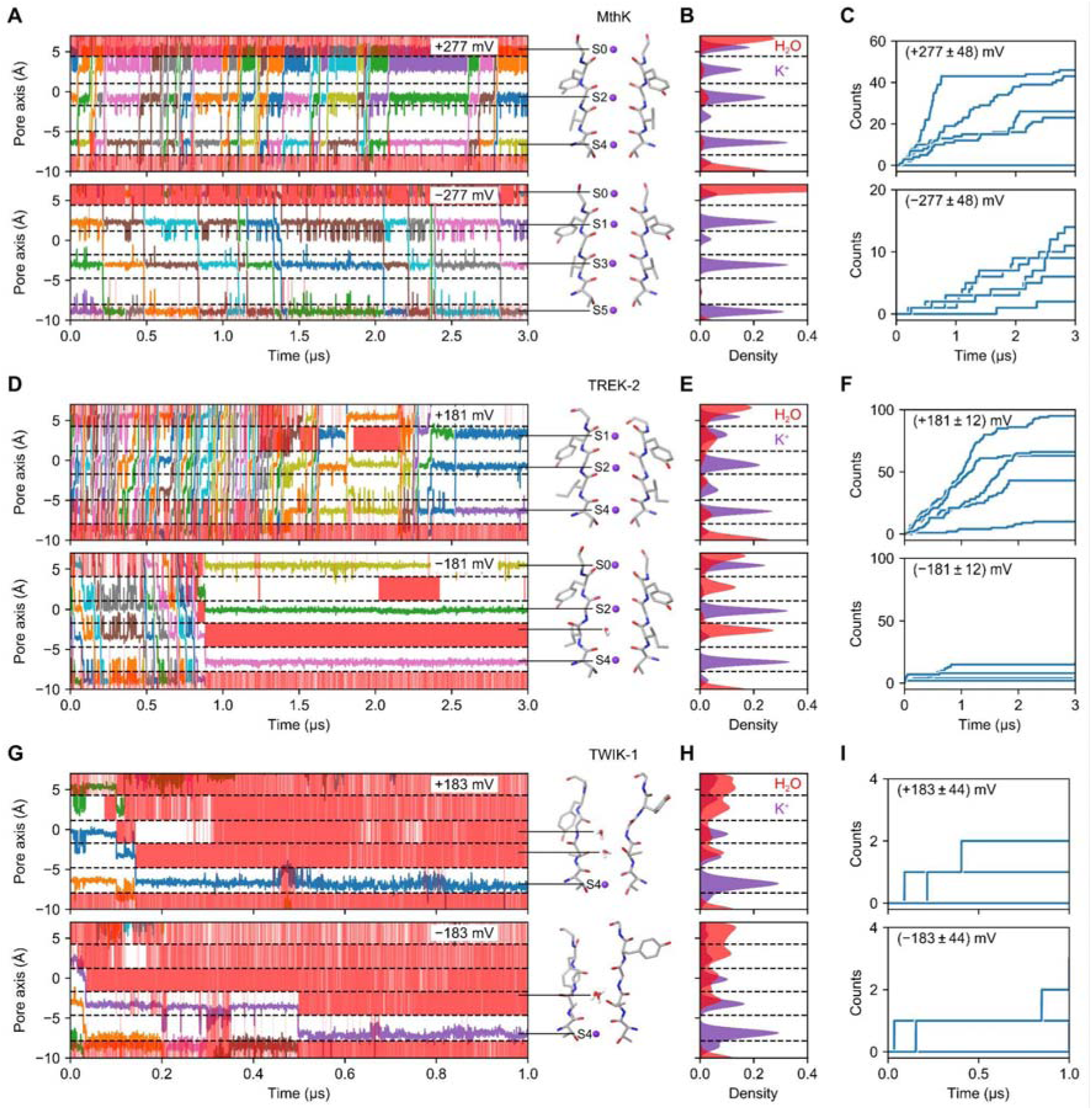
Voltage gating difference in MthK, TREK-2, and TWIK-1. (**A**, **D**, and **G**) Trajectories of K^+^ ions (different colors) along the pore axis in one run of 3 μs simulation for MthK (A) and TREK-2 (D), while 1 μs for TWIK-1 (G) under positive (upper panels) and negative (lower panels) voltages (see also Fig. S1-S3). Dashed lines indicate the center of mass of the oxygen atoms delimiting the binding sites from the respective initial structures. Red bands within the binding sites indicate the occupancy of at least one water molecule. The SF conformations are from the last frames of the respective simulations. (**B**, **E**, and **H**) K^+^ (purple) and water (red) density in the SF of MthK (B), TREK-2 (E), and TWIK-1 (H) under positive (upper panels) and negative (lower panels) voltages calculated from five simulation runs. (**C**, **F**, and **I**) Cumulative K^+^ permeation events in each simulation run for MthK (C), TREK-2 (F), and TWIK-1 (I) under positive (upper panels) and negative (lower panels) voltages. The voltage values are shown as mean ± SEM calculated from five simulation runs.

For MthK, a canonical K^+^ channel with low voltage sensitivity, we observed steady inward and outward K^+^ permeations using the same simulation setup (Fig. 2A, C, Fig. S1). Although the number of outward ion permeations was considerably higher than the inward ones, the S1-S4 sites remained largely dehydrated under both positive and negative voltages (Fig. 2B). Interestingly, under positive voltages K^+^ ions mainly occupied the S0, S1, S2, and S4 sites, while S1, S3, and S5 were primarily occupied under negative voltages (Fig. 2B).

In strong contrast to the TREK-2 and MthK simulations, TWIK-1 exhibited very low outward and inward K^+^ permeations, accompanied by the replacement of K^+^ ions by water and an accumulation of water molecules at the S1-S3 sites (Fig. 2G, I, Fig. S3). Similar K^+^ and water occupancies were observed between simulations under positive and negative voltages (Fig. 2H). This result is in good agreement with the low intrinsic activity of TWIK-1 and also aligns with our previous TWIK-1 simulations at much shorter timescale (*32*) using the Amber99SB force field (*36*). Overall, the comparison of simulations of these three types of K^+^ channels, which exhibit markedly different conductance and voltage response, suggests that the observed differences in outward and inward permeations for TREK-2 are intrinsic to the SF and might directly relate to its voltage gating behavior.

We further investigated the impact of voltage on the dynamics of the SF residues by analyzing the time evolution and distribution of backbone dihedral angles across the investigated channels (Fig. 3A, Fig. S4-S6). A quantitative measure of SF stability was obtained by integrating the density distribution of the ψ angle for each SF residue, as shown in Fig. 3A. In this analysis, values close to 1 indicate that the backbone conformation of the respective residue remained close to the experimentally determined structure throughout the simulations, while values close to 0 suggest significant deviation and thus instability of the respective residue. This analysis indicated that the SF of MthK was highly stable under both positive and negative voltages (Fig. 3A, Fig. S4), contributing to a high K^+^ occupancy in the filter. In contrast, TWIK-1 exhibited a highly dynamic SF (Fig. 3A, Fig. S6), leading to reduced K^+^ binding in the filter and consequently low K^+^ permeations under both positive and negative voltages.

**Figure 3.**
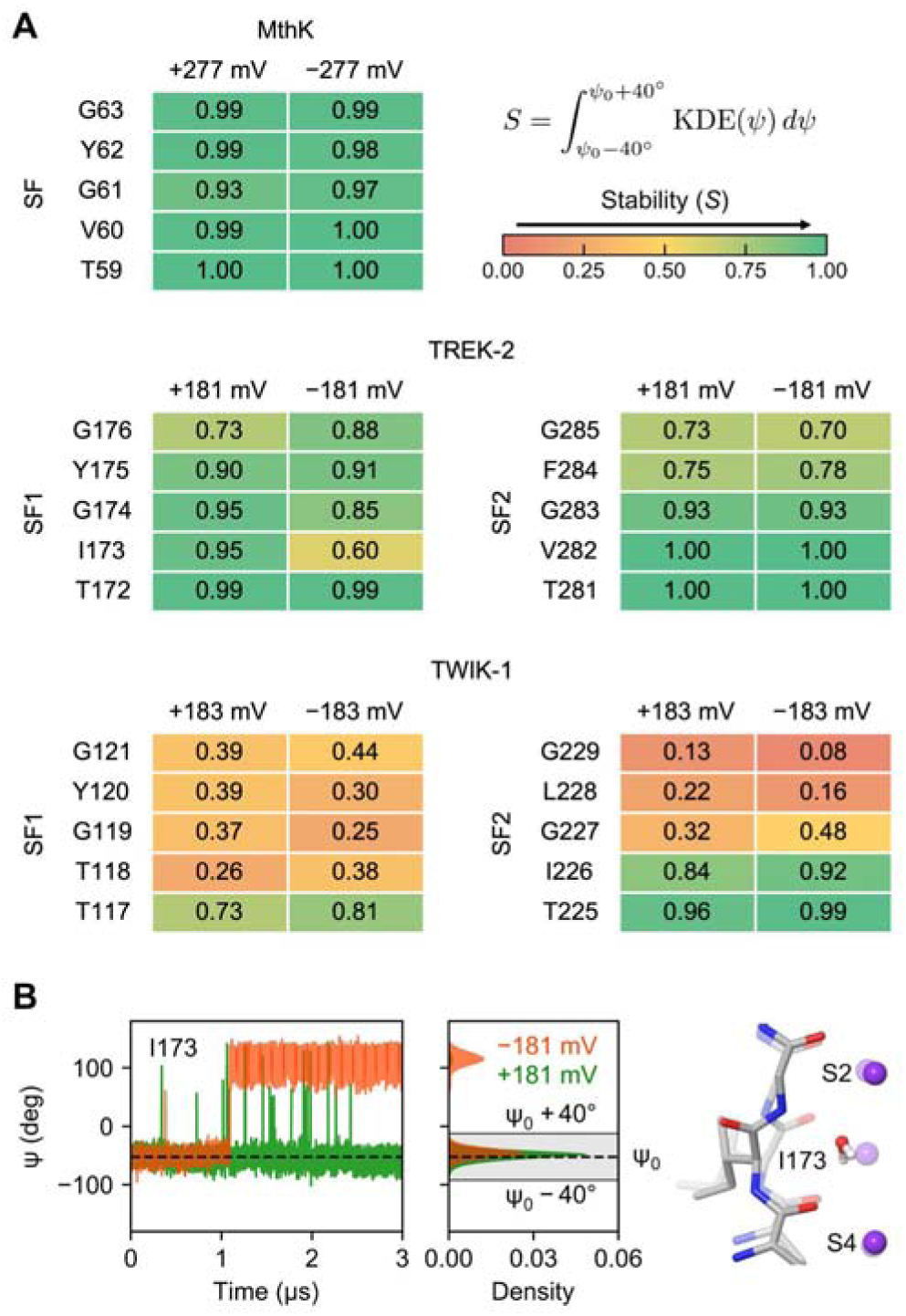
SF stability in MthK, TREK-2, and TWIK-1. (**A**) Stability of the SF residues of MthK, TREK-2, and TWIK-1 quantified by calculating the integral of the Gaussian kernel density estimation (KDE) function of the ψ angle on an interval [ψ_0_ – 40°, ψ_0_ + 40°], where ψ_0_ is the angle from the respective initial structure. KDE of the ψ angle of each SF residue was obtained from five runs of 3 μs simulations for TREK-2 and MthK, while 1 μs for TWIK-1. (**B**) The ψ angle of I173 of TREK-2 over time from one simulation run (see also Fig. S5) and its KDE calculated from five simulation runs under positive (green) and negative (orange) voltages. Dashed lines indicate ψ_0_ and the integration interval is shaded in the density plot. Examples of flipped and non-flipped (transparent) conformations of I173 are shown on the right.

Intriguingly, in contrast to MthK and TWIK-1, TREK-2 displayed non-uniform dynamics along the SF, with increased flexibility towards the extracellular side under both positive and negative voltages (Fig. 3A, Fig. S5). Notably, the carbonyls of F284 in SF2, which constitute the S1 site, frequently flipped away from the ion permeation pathway, rendering them incapable of coordinating K^+^ at S1. Consequently, this led to increased water occupancy at S1 under both positive and negative voltages (Fig. 2E). However, a marked difference in water occupancy at more inner binding sites was observed between simulations under positive and negative voltages. Due to the higher dynamics on the extracellular side, inward ion-flux facilitated water influx into the inner binding sites, which did not occur during outward ion flow. It is plausible that water influx into the inner binding sites destabilized the conductive SF conformation. For example, I173 in SF1, which forms the S3 site, exhibited significantly higher dynamics under negative voltages (Fig. 3A, B). This elevated level of dynamics correlated with increased water occupancy at S3 (Fig. 2E). Therefore, we propose that the differential conformational stability between the intracellular and extracellular sides of the SF, coupled with distinct water influx properties, underlies the origin of flux-mediated voltage sensitivity in TREK-2.

### Mutation within the SF alters water occupancy and voltage gating

Previously, we demonstrated that the mutation of threonine at the S4 site to cysteine in various K_2P_ channels abolishes voltage sensitivity (*19*). Our earlier simulations revealed significant differences in ion occupancy between the wide-type (WT) TRAAK and T103C mutant, where experimentally the channel maintained steady conductance under negative voltages (*19*). To further understand the differences between the WT and mutant channels in voltage gating at an atomistic scale, we introduced the equivalent mutation in TREK-2, replacing threonine at position T172 (equivalent to T103 in TRAAK) with cysteine in both chains, and performed five 3 μs simulation runs. Experimentally, the TREK-2 T172C mutant revealed a voltage-insensitive behavior (Fig. 4H-J). Simulations of this mutant under negative voltages showed steady inward K^+^ permeations in most of the simulation runs, with no significant water influx into the SF (Fig. 4A, C, Fig. S7). This finding is in stark contrast to the simulations of the WT and aligns with the loss of voltage sensitivity. Under positive voltages, the inner binding sites remained dehydrated, indicating a stable SF. However, we observed a strong reduction in ion conduction (Fig. S7). Why the simulations failed to show steady K^+^ permeations under positive voltages is currently unclear.

**Figure 4.**
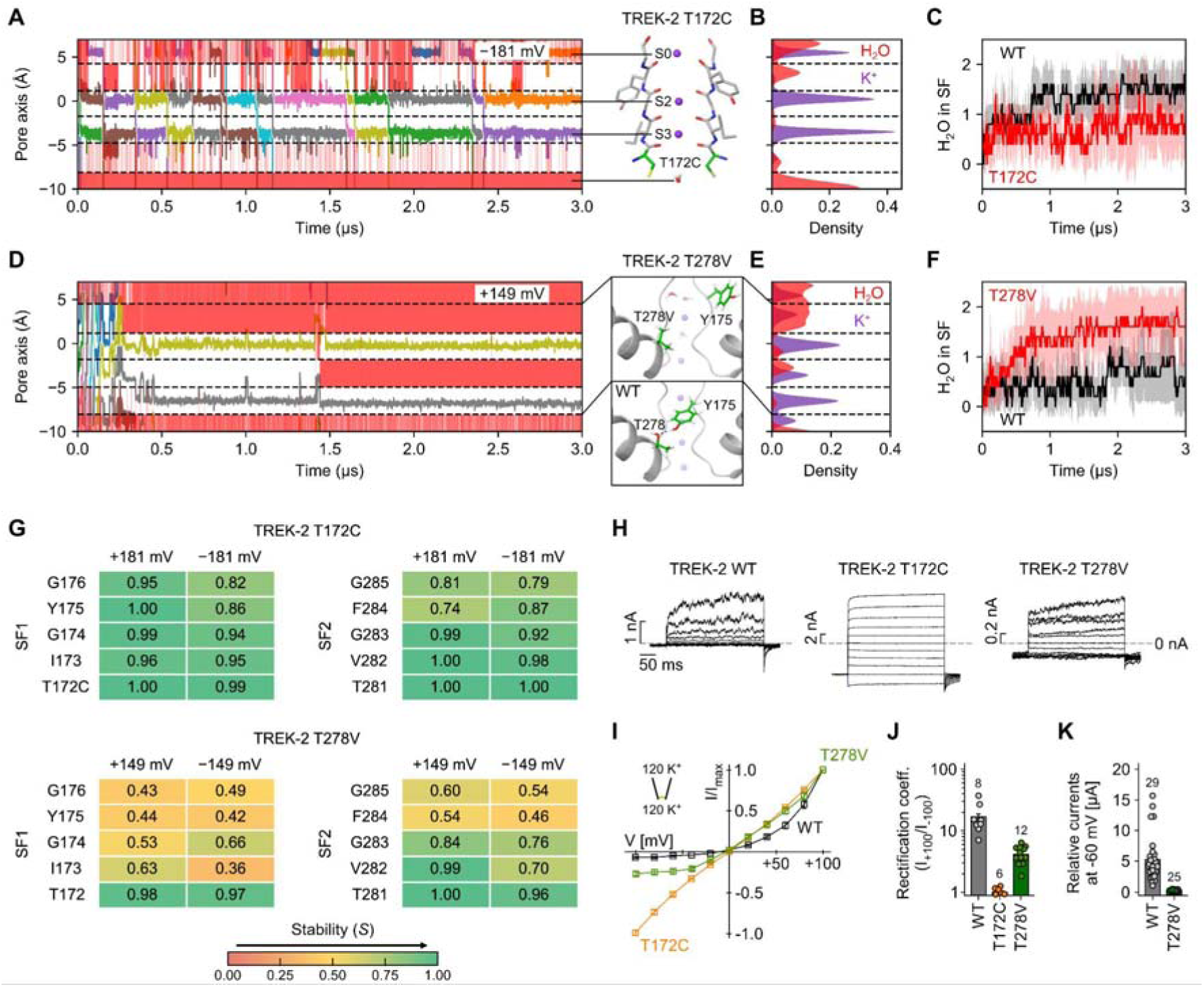
T172C and T278V filter mutants alter voltage gating in TREK-2. (**A** and **D**) Trajectories of K^+^ ions (different colors) along the pore axis in one run of 3 μs simulation for T172C mutant (A) under negative and T278V mutant (D) under positive voltages (see also Fig. S7, S13). Dashed lines indicate the center of mass of the oxygen atoms delimiting the binding sites from the respective initial structures. Red bands within the binding sites indicate the occupancy of at least one water molecule. The SF conformation and T278V mutant structure are from the last frames of the respective simulations, whereas the WT is from the initial structure of TREK-2. Key residues are highlighted in green. (**B** and **E**) K^+^ (purple) and water (red) density in the SF of T172C mutant (B) under negative and T278V mutant (E) under positive voltages calculated from five simulation runs. (**C** and **F**) Average number of water molecules in the SF of TREK-2 WT (black), T172C and T278V mutants (red) under negative (C) and positive (F) voltages (see also Fig. S9). Data shown as mean ± SEM calculated from five simulation runs. (**G**) Stability of the SF residues of the T172C and T278V mutants (see also Fig. 3A, Fig. S8, S14). (**H**) Current-voltage responses of TREK-2 WT, T172C and T278V mutants measured from –100 mV to +100 mV, with 20 mV increments in symmetrical high K^+^ [120 mM ex./120 mM int.] at pH 7.4. (**I** and **J**) I-V curves (I) and rectification coefficients (J, quotient of currents at +100 mV and –100 mV) analyzed from measurements as in (H) for the indicated TREK-2 channels. (**K**) Relative currents measured from oocytes expressing TREK-2 WT and T278V mutant in TEVC with extracellular high K^+^ [96 mM] at pH 7.4 and analyzed at –60 mV. Analyzed electrophysiological data shown as mean ± SEM from n (number of independent experiments) as indicated in the figure panel. The representative experiments were repeated with similar results.

The TREK-2 T172C mutant lacks two K^+^ coordinating oxygen atoms at the S4 site, resulting in significantly reduced K^+^ occupancy at this site compared to the WT (Fig. 4B). This observation is consistent with our previous simulations at much shorter timescale and with a different force field (*19*). Importantly, since K^+^ ions are incapable of binding to S4 in the T172C mutant, they instead preferentially occupied the S3 site during inward permeation and, thereby, antagonized water binding to the S3 site. This, in turn, stabilized I173 at S3 in its non-flipped conformation preventing SF inactivation (Fig. 4A, Fig. S7). In conclusion, our MD simulations suggest that the TREK-2 T172C mutant significantly reduces the conformational dynamics of the SF under negative voltages (Fig. 4G, Fig. S8), providing a plausible mechanism for the abolishment of voltage sensitivity in this mutant.

### MD simulations at increased temperature revealed full SF inactivation

Our five runs of 3 µs simulations of TREK-2 under negative voltages have thus far revealed a notable decrease in K^+^ occupancy and an increase of water occupancy in the SF, with an average of two K^+^ ions in the filter (Fig. S9). However, based on the gating charge analysis we previously performed on TREK/TRAAK K_2P_ channels, the fully inactivated filter should be nearly devoid of ions (*19*). This suggests that, due to the limited timescale of the simulations, the SF did not fully reach its inactivated state during these 3 µs simulations. To accelerate the filter inactivation process, we extended the simulations of TREK-2 by increasing the system temperature by 20 K (from 303 K to 323 K) to enhance the sampling. Notably, in three out of five extended 3 µs simulations at 323 K, we observed K^+^ exiting the S2 site into the extracellular solution, leaving only one K^+^ bound at S4 (Fig. 5A, Fig. S10, Mov. 5, 6). In this state, we observed substantial conformational changes at the extracellular side of the filter compared to the conductive conformation observed in the X-ray structure (Fig. S11).

**Figure 5.**
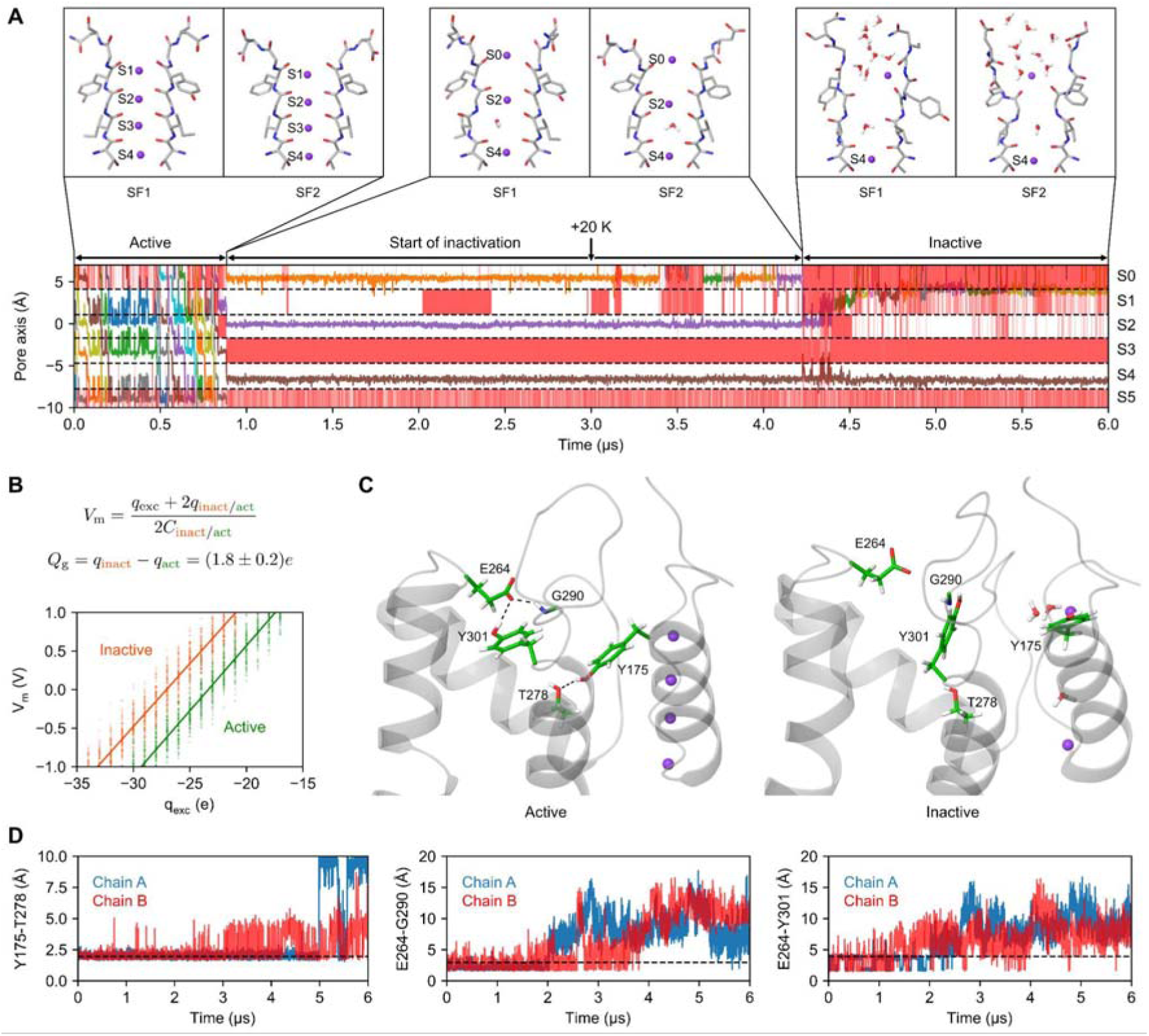
SF inactivation in TREK-2. (**A**) Trajectories of K^+^ ions (different colors) along the pore axis in one run of 6 μs simulation showing complete SF inactivation in TREK-2 under negative voltages (see also Fig. S10). Dashed lines indicate the center of mass of the oxygen atoms delimiting the binding sites from the initial structure. Red bands within the binding sites indicate the occupancy of at least one water molecule. The increase of system temperature from 303 K to 323 K at 3 μs is indicated by an arrow. Representative SF conformations correspond to three different filter states during inactivation. (**B**) Influence of the state of TREK-2 on the capacitor-like property of the system as shown in the charge titration plot, from which the gating charge value was obtained. (**C**) Three H-bonds in the active state and their disruption in the inactive state, where the side chains of key residues are highlighted in green. (**D**) Distances between donor and acceptor atoms over time in one simulation run showing the disruption of three H-bonds during SF inactivation (see also Fig. S12). Dashed lines indicate the H-bond lengths from the active state.

To validate the inactivated state obtained from the MD simulations performed with elevated temperature, we conducted a computational gating charge analysis using the CompEL setup (*37*). This analysis determines the change in the electrical capacitance of the channel during its conformational change. In our case, the active and inactive states of TREK-2 have different contributions to the capacitor charge, and this difference is used to derive the gating charge. For the active state, we used the initial structure of TREK-2 with four bound K^+^ ions in the SF. For the inactive state, we used the structure obtained from the simulation at elevated temperature, which suggested an inactivated SF containing only one K^+^ bound at S4 (Fig. 5A). The gating charge (e_0_) was calculated as 1.8 ± 0.2 (Fig. 5B). This result agrees well with electrophysiological gating charge measurements that determined an equivalent gating charge between 1.8 and 2.6 e_0_ for voltage gating in K_2P_ channels (*19*).

### Disruption of three H-bonds during SF inactivation

By analyzing the MD trajectories of SF inactivation, we observed the disruption of three different H-bonds in the extended filter gate of TREK-2 (Fig. 5C, D, Fig. S12). Two H-bonds are part of the “TM3 glutamate network”, specifically between E264 and residues G290 and Y301 (Fig. 5C). The TM3 glutamate network has previously been shown to stabilize the unusually long loop connecting SF2 with TM4 (*5*). The other H-bond exists between the SF residue Y175 and T278 located in the pore helix. This H-bond stabilizes the active conformation of the filter gate by reducing its dynamics at the extracellular side.

To evaluate the importance of the Y175-T278 H-bond for SF stability, we studied the TREK-2 T278V mutant. Whole-cell measurements of this mutant channel revealed a significant reduction in current (Fig. 4H, K), although the channel remained voltage-gated (Fig. 4H-J). We also conducted ion permeation simulations of the TREK-2 T278V mutant. These simulations under positive voltages showed a significantly reduced number of K^+^ permeation events and a highly dynamic SF compared to the WT (Fig. 4D-G, Fig. S13, S14). The increased dynamics of the SF under both positive and negative voltages allowed water molecules to permeate and occupy the S3 site, effectively halting K^+^ permeations in most simulation runs (Fig. 4D, Fig. S13), in agreement with the electrophysiological phenotype.

### The role of extracellular water molecules for SF inactivation

The MD simulations predict that extracellular water molecules regulate conformational changes in the SF that underlie ion-flux gating in TREK-2. Specifically, water entering into the S3 site leads to SF inactivation. However, water can only enter the filter from the extracellular side, while the intracellular SF entrance appears to be watertight. To test this concept, we reduced the availability of water by adding a high concentration of sucrose (2 M) to either the extracellular or intracellular solution (Fig. 6A). The strong prediction is that ion-flux activation arising from K^+^ entering the inactivated filter from the intracellular side should remain unaffected by sucrose. In contrast, ion-flux inactivation—if it indeed arises from extracellular water entering the SF—should be slowed due to the complexation of water with sucrose.

**Figure 6.**
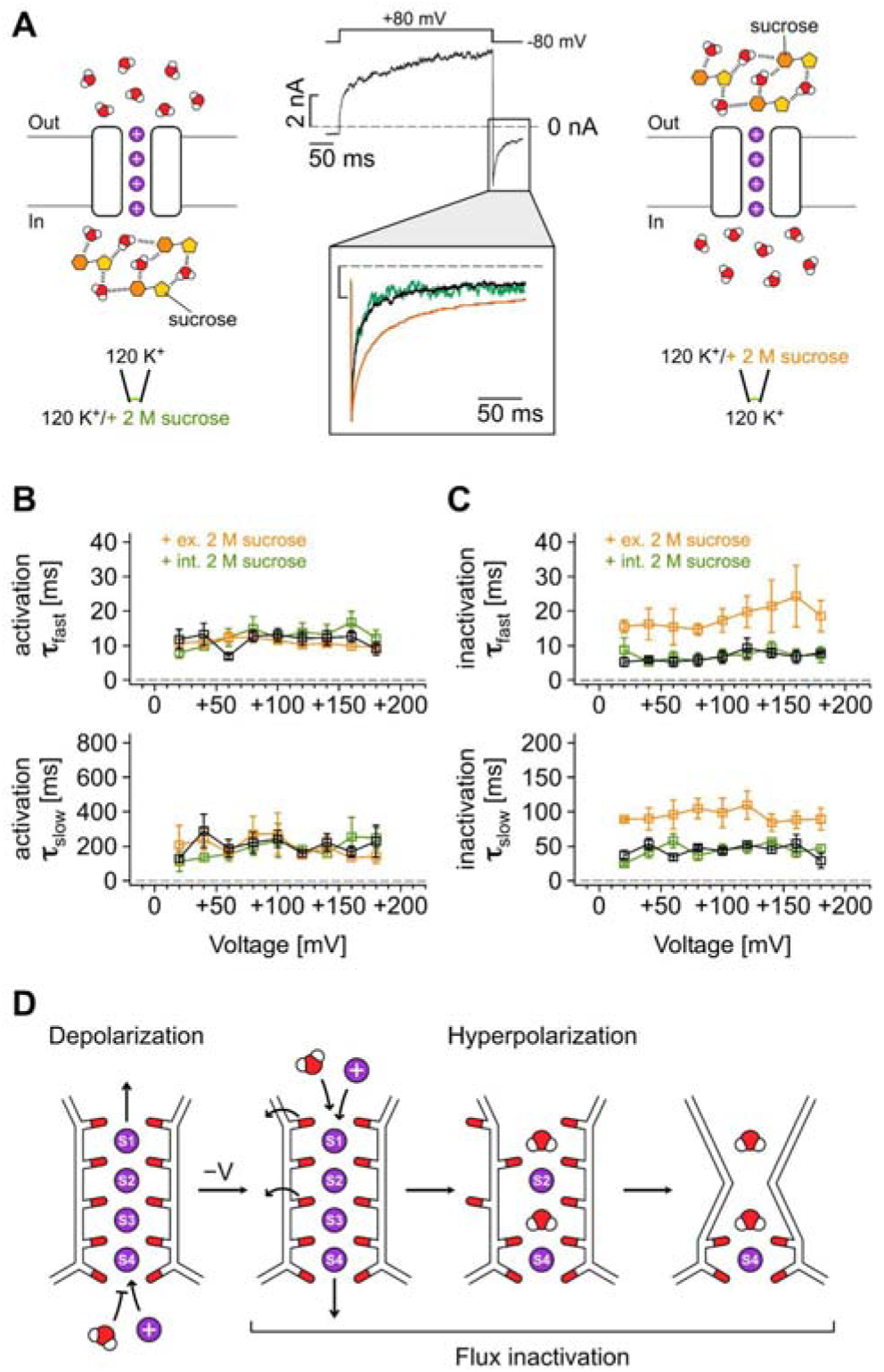
Non-canonical voltage gating in TREK K_2P_ channels. (**A**) Representative current trace of TREK-2 recorded in symmetrical high K^+^ as indicated at pH 7.4 showing time– and voltage-dependent activation at 300 ms depolarizing +80 mV and a tail current amplitude at hyperpolarizing –80 mV potential. Zoom-in shows an overlay of recordings without (black) and with extracellular (orange) or intracellular (green) 2 M sucrose. Schematic illustrations highlighting the effect of sucrose on water molecules at either side are shown. (**B** and **C**) Analysis of fast (__fast_) and slow (__slow_) time constants of activation (B) and inactivation (C) from measurements as in (A) for the indicated voltages in the absence (black) and presence of 2 M sucrose at the extracellular (orange) or intracellular (green) side of the membrane. Analyzed electrophysiological data shown as mean ± SEM from n (number of independent experiments) as indicated in the figure panel. (**D**) Schematic illustration of the atomistic mechanism of non-canonical voltage gating in TREK K_2P_ channels.

To evaluate this, we induced ion-flux gating in TREK-2 by applying depolarizing voltage steps (from +20 to +180 mV) to induce outward flux activation, followed by hyperpolarization (to –80 mV) to induce inward flux inactivation (apparent as tail currents). These experiments were performed using symmetrical K^+^ solutions in inside-out patches. In TREK-2, both ion-flux activation and inactivation displayed a biexponential time course. Although the exact cause of the biexponential kinetics remains unclear, this behavior is not surprising, as both activation and inactivation gating involve multiple reaction steps (i.e. several K^+^/water binding and movement steps, followed by SF activation/inactivation). Intriguingly, sucrose had a selective effect on the ion-flux gating transitions: voltage activation remained unaffected by sucrose, regardless of whether it was present extracellularly or intracellularly (Fig. 6B). In contrast, the time course of inactivation was markedly slowed in the presence of extracellular sucrose but not intracellular sucrose (Fig. 6C). Specifically, the fast component of inactivation was slowed by a factor of 5, while the slow component was slowed by a factor of 2. These findings suggest that sucrose selectively affects the SF gating transition that depends on the availability of extracellular water (i.e. inward flux inactivation). This result supports the MD predictions, which propose that the SF allows water entry from the extracellular side to terminate inward permeation, leading to filter inactivation, while preventing water entry from the intracellular side. This mechanism is illustrated in Fig. 6D.

## Discussions

In the current study, we conducted large-scale atomistic MD simulations of ion permeation in three types of K^+^ channels to investigate the atomistic basis of ion-flux gating in K_2P_ channels. We selected TREK-2, TWIK-1, and MthK channels, as TREK-2 exhibits prominent ion-flux gating, while TWIK-1 and MthK lack this property. Our aim was to explore whether these differences arise from specific properties of the SF.

Our findings highlight the presence of water molecules in the SF as the key factor underlying ion-flux gating, with the stability of the filter being the primary determinant for water entry. The SF of MthK was very stable under both positive and negative voltages, devoid of water, and capable of steadily conducting outward and inward ion permeation. This behavior aligns with the high conductance and open probability characteristic of this channel. However, our simulations using the conventional CHARMM36m force field parameters failed to reproduce the inward-rectifying property of MthK. Notably, a recent study employing charge-scaled K^+^ parameters successfully replicated this characteristic (*38*). In stark contrast to MthK, the SF of TWIK-1 was highly dynamic, allowing water molecules to enter the filter under both positive and negative voltages. The resulting high water occupancy within the SF significantly slowed down ion permeation, consistent with the very low intrinsic open probability of TWIK-1.

The SF in TREK-2, however, exhibited an intriguing asymmetric stability difference between the intracellular and extracellular sides. The intracellular side of the SF was stable, devoid of water, and capable of steadily conducting outward currents. In contrast, the extracellular side of the SF was more dynamic, allowing water to enter the filter and occupy the S1 site. The water then accumulated at the S3 site, causing ion permeation stop and inducing backbone carbonyl flip at S3, which marked the first step of SF inactivation. This event was followed by the unbinding of three K^+^ ions, ultimately leading to the final inactivated state, as observed in MD simulations performed at elevated temperature. This finding aligns well with the previously proposed ion-flux gating mechanism in TREK-2 and other K_2P_ channels, based on electrophysiology measurements. A hallmark of this mechanism is the steep dependence of the SF open probability (*P_O_*) on the electrochemical gradient. Thus, K_2P_ channels are not inherently voltage-sensitive but are instead sensitive to the direction of ion-flux. Our current MD simulations provide an atomistic understanding of this ion-flux dependence. At potentials increasingly positive relative to the reversal potential, the likelihood of K^+^ ions moving outward through the SF and displacing water from the filter increases. This reduces the probability of water occupying the S1 and S3 sites, thereby preventing filter inactivation. Conversely, at potentials negative to the reversal potential, the instability of the S1 site to coordinate K^+^ becomes more pronounced. At strong negative potentials, the S1 site becomes almost exclusively occupied by water, promoting water movement to the S3 site and triggering SF inactivation. Therefore, the ion-flux gating properties of TREK-2 can be described, figuratively, as a competition between extracellular water and intracellular K^+^ for occupancy of the S3 site. Water binding at this site induces SF inactivation, while K^+^ binding prevents or reverses inactivation. This dynamic ultimately determines the *P_O_*of the SF.

In this study, we confirmed the critical role of extracellular water in SF inactivation through electrophysiology measurements by comparing the effects of adding sucrose to the intracellular versus extracellular solution. Sucrose has previously been used as a biophysical probe to study SF size, conductance properties, and filter inactivation in various K^+^ channels (*39*). For instance, a previous study on the bacterial K^+^ channel KcsA demonstrated that reducing buried water molecules behind the SF, enabled by adding extracellular sucrose, accelerates filter inactivation (*40*). In our study, we found that introducing sucrose to both the intracellular and extracellular sides had no effect in voltage activation of TREK-2. However, the inactivation kinetics were markedly slowed when extracellular sucrose was added, an effect that was not observed with intracellular sucrose addition.

Furthermore, we noted previous MD simulations investigating mechanosensitivity of TREK-2 under membrane stretch also revealed a correlation between conductance and the stability of the S3 site (*41*). Additionally, a recent MD study exploring the mechanistic effects of two TREK-1 activators binding behind the SF demonstrated that these ligands reduce carbonyl flipping at S3, further stabilizing the threonine at S4 (*42*). Together with our findings, these MD studies, which focus on different biophysical aspects of TREK K_2P_ channels, collectively underscore the critical role of the stability of the S3 site in channel inactivation. This stability is closely associated with water occupancy and represents a key determinant in the SF inactivation process.

Furthermore, based on a comparative analysis of the TREK-2, TWIK-1, and MthK channels, we propose that structural instability at the extracellular side of the SF is another key contributor to ion-flux gating in TREK K_2P_ channels. This instability facilitates water influx, which leads to SF inactivation (Fig. 6D). Additional experimental evidence of the high dynamics at the extracellular side is provided by the structures of TREK-1 determined at low K^+^ concentrations (*5*), and the TASK-2 and TASK-3 structures resolved at low pH levels (*15*, *16*). In TASK-2, conformational instability at the extracellular side leads to the loss of K^+^ binding at S1 ((*15*, *16*)), while both S1 and S2 sites are abolished in the filter-inactivated state of TREK-1 and TASK-3 structures (*5*, *16*). Moreover, in a recent study, using a combination of electrophysiology and MD simulations, we demonstrated that structural changes at the extracellular side of the SF are coupled with conformational dynamics of the TM4 helix and the proximal C-terminus, which senses external stimuli such as phosphorylation (*43*). Notably, the MD simulations in this previous study were performed with the Amber99SB force field, yet similar conformational rearrangements at the extracellular side were observed. Taken together, we propose that conformational instability at the extracellular side of the SF plays a central role in the control of K_2P_ channels by external stimuli such as voltage, pH, or phosphorylation.

In good agreement with MD simulations of other K^+^ channels performed using various force fields (*44–46*), the ion-mediated knock-on mechanism also dominated the outward K^+^ permeation process in our TREK-2 simulations. The water influx observed during inward K^+^ flow led to SF inactivation. Therefore, our findings reveal that the ion-mediated knock-on is not only an efficient mechanism for K^+^ conduction, but also plays a critical role in maintaining the conductive state of TREK-2.

In contrast to other reported MD works so far, we showed that under full SF inactivation only one K^+^ remains bound at S4. The computed gating charge based on this inactive state of the SF aligns closely with previously measured experimental values. This ion configuration differs from the X-ray structures of TREK-1 determined at low K^+^ concentrations, which revealed ion binding at both S3 and S4. Furthermore, we noted that during complete inactivation, the SF undergoes large conformational changes at the extracellular side. Similarly, the X-ray structure of TREK-1 at low K^+^ concentration revealed pinched SF1 and dilated SF2 at the extracellular side (*5*). The conformational change in the SF during inactivation also leads to the disruption of two H-bonds in the TM3 glutamate network together with the breakage of H-bond between T278 and Y175. This finding suggests that during the SF inactivation of TREK-2, conformational instability can propagate from the filter to the connecting loop regions.

We also performed MD simulations of TRAAK (PDB: 4I9W) and observed a similar tendency for increased water occupation at S1 and S3 under negative voltages, although the inactivation kinetics appears to be slower compared to TREK-2 (Fig. S15, S16). The K_2P_ channels, excluding the members of the TWIK subfamily, show similar voltage gating characteristics and have the sequence of T(I/V)G(Y/F)G in SF1 and T(I/V)GFG in SF2. Therefore, the proposed atomistic voltage gating mechanism of the TREK subfamily may extend to other K_2P_ channels, although systematic studies using similar approaches combining MD simulations and electrophysiological measurements are needed.

Due to the central role of SF flexibility in the gating of K_2P_ channels, the voltage sensitivity and conductance properties of TREK K_2P_ channels are highly influenced by mutations within and around the SF. Furthermore, it should be noted that voltage gating in TREK channels can also be modulated pharmacologically. Small molecule activators, such as BL-1249, that bind below the SF, abolish voltage gating behavior in TREK by stabilizing the filter and reducing its dynamics through alterations of ion occupancy in the pore and consequently the filter gate (*23*).

In conclusion, we propose an atomistic mechanism for non-canonical voltage gating in TREK K_2P_ channels (Fig. 6D), which is enabled by water influx under hyperpolarization that initiates SF inactivation. This mechanism is supported by a systematic comparison of three K^+^ channels with different voltage gating behaviors, as well as by electrophysiological data and MD simulations of two TREK-2 mutants. The most compelling validation of the key role of extracellular water in channel inactivation under hyperpolarization comes from the electrophysiological measurements using sucrose (Fig. 6A-C). The voltage gating mechanism in TREK K_2P_ channels is fundamentally different from those in other ion channels and could pave the way for understanding and engineering voltage sensitivity in other ion channels and membrane proteins.

## Methods

### MD simulations

Simulation systems were prepared by CHARMM-GUI (*47*) with PDB: 3LDC (MthK) (*48*), PDB: 4BW5 (TREK-2) (*7*), PDB: 7SK0 (TWIK-1) (*3*), and PDB: 4I9W (TRAAK) (*9*) as initial structures. All residues were assigned their standard protonation states at pH 7. N and C termini were capped with acetyl and methylamide groups, respectively. Structures were embedded in POPC (1-palmitoyl-2-oleoyl-sn-glycero-3-phosphocholine) and solvated in 600 mM KCl. CHARMM36m (*35*) was used as a force field and TIP3P (*49*) was used as a water model. Simulations were initiated with the channels containing four K^+^ ions bound at S1-S4 sites and run using GROMACS 2021 (*50*). Systems were energy minimized for 5000 steps followed by a multistep equilibration in which protein and lipid restraints were gradually reduced over 10 ns. Next, double bilayer systems were built following the CompEL protocol (*34*) with the charge imbalance set to 2e (to generate the membrane potential) and equilibrated for 20 ns without restraints (Fig. S17). A deterministic protocol within the CompEL framework was employed to maintain ion imbalance throughout the simulations. Production simulations used 2 fs time step, Parrinello-Rahman barostat (*51*) with semi-isotropic pressure control at 1 atm, and a v-rescale thermostat (*52*) set to 303.15 K. Nonbonded interactions were cut off at 12 Å with force-switching between 10 and 12 Å, long-range electrostatics were calculated with particle mesh Ewald (*53*), and hydrogens were constrained with the LINCS algorithm (*54*). Mutations to TREK-2 (PDB: 4BW5) were introduced using CHARMM-GUI and simulated using the same methodology as for WT. In extended TREK-2 simulations the system temperature was set to 323.15 K after 3 μs.

### Analysis of the MD data

Trajectories were saved every 50 or 100 ps and analyzed using MDAnalysis (*55*) and NumPy (*56*) Python libraries. K^+^ ions were tracked within a cylindrical zone of radius 15 Å and height 40 Å centered on the SF. Water molecules were tracked within a cylindrical zone of radius 5 Å and height 22 Å centered on the SF. A permeation event was recorded when an ion bound in the SF crossed the midplane of the filter perpendicular to the pore axis last time. Ion and water density within the S0-S5 sites were obtained using Gaussian KDE from SciPy (*57*) with default parameters. Dihedral angles were calculated using *gmx rama* from GROMACS and the ψ angle density of each SF residue was obtained using Gaussian KDE from SciPy with default parameters. Distances between atoms were calculated using *gmx distance*. The electrostatic potential along the pore axis was computed using *gmx potential* and the voltage was calculated as the difference in electric potential between two sides of the bilayer. Plots were generated using Matplotlib and protein visualizations were made with PyMOL and Maestro. Movies were created using VMD.

### Gating charge analysis

An antiparallel CompEL system of the active TREK-2 state with four K^+^ ions bound at the S1-S4 sites was built and equilibrated for 20 ns with position restraints with a force constant of 1000 kJ mol^-1^ nm^-2^ applied to backbone atoms and four K^+^ ions in the SF. For the inactive TREK-2 state, a new system was prepared by CHARMM-GUI using the inactivated structure with only one K^+^ bound at S4 extracted from one extended simulation. The system with inactive TREK-2 state was energy minimized and equilibrated, after which an antiparallel CompEL system was built and equilibrated for 20 ns with position restraints with a force constant of 1000 kJ mol^-1^ nm^-2^ applied to backbone atoms and one K^+^ at S4. After preparing the antiparallel CompEL systems with the active and inactive TREK-2 states, 100 ns long MD simulations were run at various ionic charge imbalances to obtain the relationship between the membrane potential and the excess charge. Position restraints were kept during production runs to keep the system stable at high voltages. All other MD parameters were the same as described in the “MD simulations” subsection. From the linear regression between the membrane potential and the excess charge, the charge contributions of the active and inactive states to the capacitor charge were obtained. Difference in charge contributions of the active and inactive states provided the gating charge.

### Molecular biology

In this study the coding sequences of human K_2P_1.1 TWIK-1 (Genbank accession number: NM_002245) and human K_2P_10.1 TREK-2 (NM_021161) were used. For K^+^ channel constructs expressed in oocytes the respective K^+^ channel subtype coding sequences were subcloned into the oocyte expression vector pBF or the dual-purpose vector pFAW which can be used for mammalian cell expression as well and verified by sequencing. All mutant channels were obtained by site-directed mutagenesis with custom oligonucleotides. To increase plasma membrane expression, measurements of TWIK-1 channels were done using channels with mutated retention motif (TWIK-1 I293A/I294A (TWIK-1*)). Vector DNA was linearized with NheI or MluI and cRNA synthesized *in vitro* using the SP6 or T7 AmpliCap Max High Yield Message Maker Kit (Cellscript, USA) or HiScribe^®^ T7 ARCA mRNA Kit (New England Biolabs) and stored at –20 °C (for frequent use) and –80 °C (for long term storage).

### Animals and cell preparation

The performed investigation conforms to the guide for the Care and Use of laboratory Animals (NIH Publication 85-23). For this study, we used female *Xenopus laevis* animals (10) that were accommodated at the animal breeding facility of Kiel University to isolate oocytes. Experiments using *Xenopus* toads are approved by the local ethics commission. *Xenopus laevis* ovarian lobes were surgically removed from tricaine anesthetized adult female frogs and treated with type II collagenase (2 mg/ml, Sigma-Aldrich/Merck, Germany) in OR2 solution containing (in mM): 82.5 NaCl, 2 KCl, 1 MgCl_2_, 5 HEPES (pH 7.4 adjusted with (NaOH/HCl) for 2 h prior to manual defolliculation. Isolated oocytes were stored at 16.9 °C in ND96 recording solution (in mM): 96 NaCl, 2 KCl, 1.8 CaCl_2_, 1 MgCl_2_, 5 HEPES (pH 7.5 adjusted with NaOH/HCl) supplemented with Na-pyruvate (275 mg/l), theophylline (90 mg/l), and gentamicin (50 mg/l). Oocytes were injected with channel-specific cRNA (0.5 – 1 µg*µl^-1^) and incubated at 16.9 °C for 1-7 days prior to the experimental use.

### Electrophysiological recordings in oocytes

#### Two-electrode voltage-clamp (TEVC) measurements

Whole-cell recordings were performed under voltage-clamp conditions using the TEVC technique and the oocyte system. Oocytes were injected with 50 ng of cRNA for WT or T278V mutant TREK-2 and incubated for 24 hrs at 16.9 °C. TEVC measurements were performed at room temperature (21 – 22 °C) with an HEKA multi electrode clamp amplifier iTEV 90 and PatchMaster software (HEKA electronics, Germany). Microelectrodes were fabricated from glass pipettes, back-filled with 3 M KCl, and had a resistance of 0.2 – 1.0 MΩ.

#### Inside-out patch-clamp measurements

Excised-patch recordings in inside-out configuration under voltage-clamp conditions were performed at room temperature (21 – 22°C). Patch pipettes were made from thick-walled borosilicate glass GB 200TF-8P (Science Products, Germany), had resistances of 0.2 – 0.5 MΩ (tip diameter of 10 – 25 µm) and filled with a pipette solution (in mM): 120 KCl, 10 HEPES and 3.6 CaCl_2_ (pH 7.4 adjusted with KOH/HCl) or 120 KCl, 10 Hepes, 3.6 CaCl_2_ and 2 M sucrose (pH 7.4 adjusted with KOH/HCl). Intracellular bath solutions were applied to the cytoplasmic side of excised patches via gravity flow multi-barrel pipette system. Intracellular solution had the following composition (in mM): 120 KCl, 10 HEPES, 2 EGTA and 1 Pyrophosphate (pH 7.4 adjusted with KOH/HCl). For control experiments the intracellular bath solution contained 2 M sucrose. Currents were recorded with an EPC10 amplifier and PatchMaster software (HEKA electronics, Germany) and sampled at 10 kHz or higher and filtered with 3 kHz (–3 dB) or higher as appropriate for sampling rate.

### Data acquisition and statistical analysis

Data analysis and statistics were done using Fitmaster (HEKA electronics, version: v2×73.5, Germany), Microsoft Excel 2021 (Microsoft Corporation, USA) and Igor Pro 9 software (WaveMetrics Inc., USA). Channel currents were recorded and analyzed at voltages defined in the respective figure legend or with a voltage protocol as indicated in the respective figure. For analysis of activation and inactivation time constants (_) current traces were fitted with a double-exponential equation as depicted in 1:

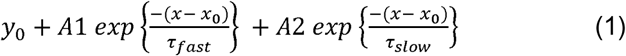

Conductance-voltage (G-V) relationships were determined from tail currents recorded at a hyperpolarizing potential of –80 mV after 300 ms depolarizing steps as indicated in the respective figure legend. Data were analyzed with a single Boltzmann fit following equation 2:

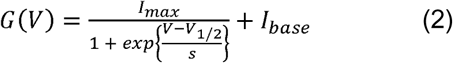

where *V*_1/2_ represents the voltage of half-maximal activation, *s* is the slope factor, and *I*_max_ and *I*_base_ represent the upper and lower asymptotes. Under appropriate experimental conditions, you can use slope to calculate the valence (charge) of the ion moving across the channel. Slope equals *R * T / z * F* where *R* is the universal gas constant, *T* is temperature in K, *F* is the Faraday constant, and *z* is the valence.

Throughout the manuscript all values are represented as mean ± SEM with n indicating the number of individual executed recordings of single patches or oocytes. Data from independent measurements (biological replicates) were normalized and fitted independently to facilitate averaging. Zero current levels are shown as dashed lines in all figures. Image processing and figure design was done using Igor Pro 9 (64 bit) (WaveMetrics, Inc., USA) and Canvas X Draw (Version 20 Build 544) (ACD Systems, Canada).

## Data availability

Input data for MD simulations and results supporting the findings and conclusions of this paper are available at Zenodo (https://zenodo.org/records/13844321?preview=1&token=eyJhbGciOiJIUzUxMiJ9.eyJpZCI6Ijc3NjIyZjM3LTJjOTEtNDZhMC1iMmEzLWU1NWNhYzEwOWI5MSIsImRhdGEiOnt9LCJyYW5kb20iOiJmMmM3Nzk2MDBjMDUzODZlZmVkNWZiZjRhZjk0YjNlZCJ9.KBruLo_8FASfIjaX7hJf4QZTt8k7vNJSLUU6mmBmaw6Kp7TEOaYNqdlcwJU-45mz6QC3I5PfygTyYytHcYe9Ig).

## Acknowledgments

We thank Denis Qoraj and Adam Lange for helpful discussions. This work was funded by the Deutsche Forschungsgemeinschaft through the Research Unit 2518 DynIon, project P01 to M.S. and T.B. and project P03 to H.S., and Leibniz Collaborative Excellence to H.S. Y.K.A. thanks Studienstiftung des Abgeordnetenhauses von Berlin for providing a research scholarship. The authors gratefully acknowledge the computing time made available to them on the high-performance computer “Lise” at the NHR Center NHR@ZIB. This center is jointly supported by the Federal Ministry of Education and Research and the state governments participating in the NHR (www.nhr-verein.de/unsere-partner).

## Author contributions

Conceptualization: HS, YKA, MS, TB Investigation: YKA, CCD, MS Formal analysis: YKA, SH, MS, TB, HS Funding acquisition: HS, MS, TB Visualization: YKA, SH, MS Writing—original draft: YKA, HS Writing—review & editing: YKA, HS, MS, TB

