## Supplementary information for "Atomistic Mechanism of Non-Canonical Voltage Gating in TREK K_2P_ Channels"

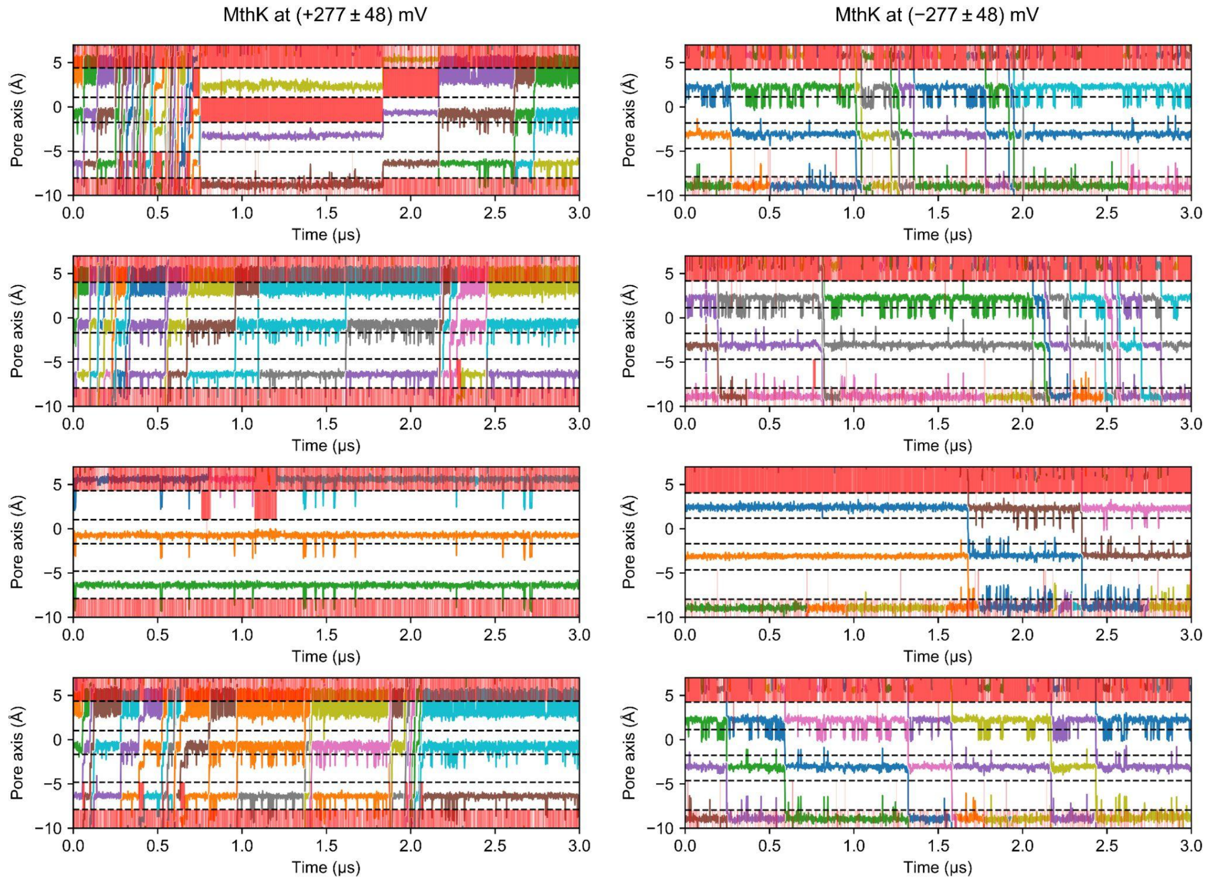


**Figure S1. Trajectories of $K^+$ ions along the pore axis in MthK.** Dashed lines indicate the center of mass of the oxygen atoms delimiting the binding sites from the initial structure. Red bands within the binding sites indicate the occupancy of at least one water molecule. The voltage values are shown as mean ± SEM calculated from five 3 μs simulation runs at 303.15 K using CHARMM36m force field. Results from one simulation are shown in Fig. 2A.

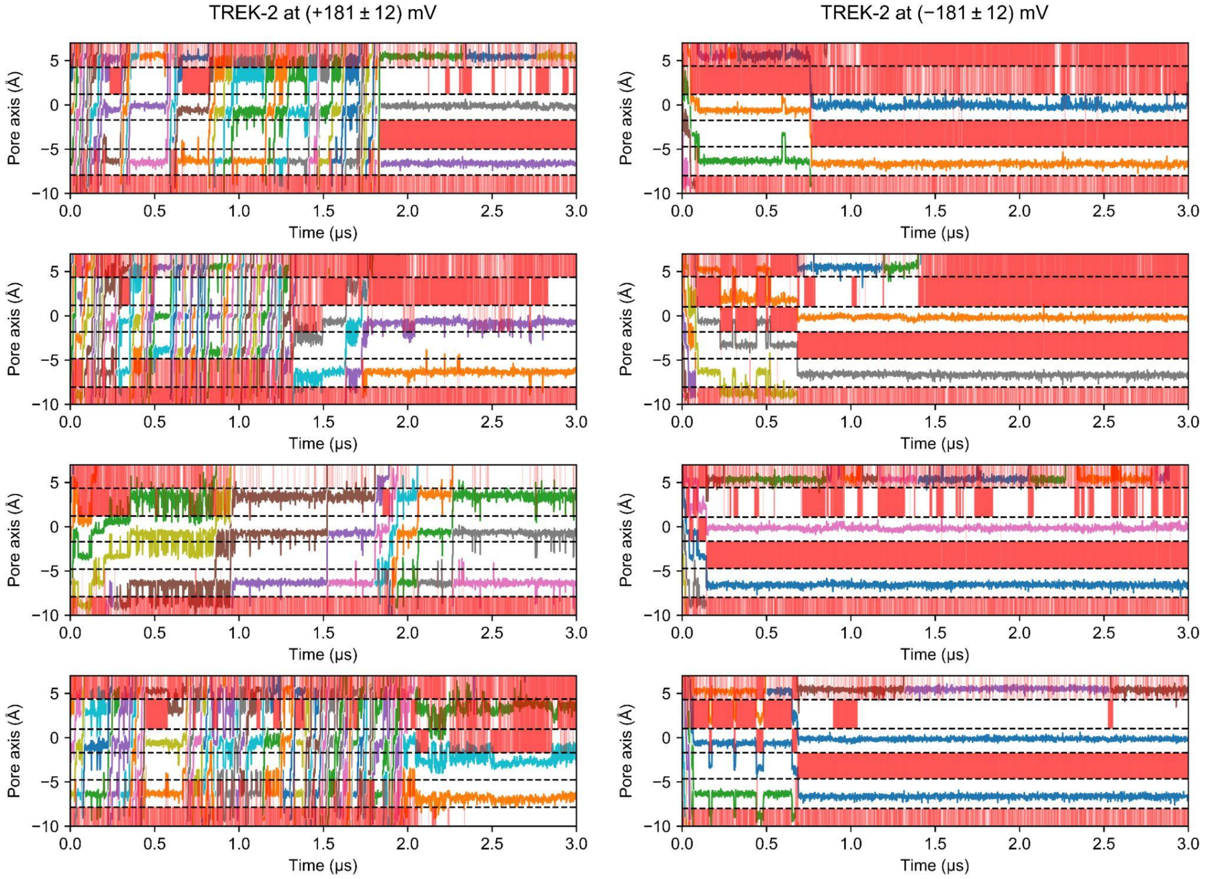


**Figure S2. Trajectories of $K^+$ ions along the pore axis in TREK-2.** Dashed lines indicate the center of mass of the oxygen atoms delimiting the binding sites from the initial structure. Red bands within the binding sites indicate the occupancy of at least one water molecule. The voltage values are shown as mean ± SEM calculated from five 3 μs simulation runs at 303.15 K using CHARMM36m force field. Results from one simulation are shown in Fig. 2D.

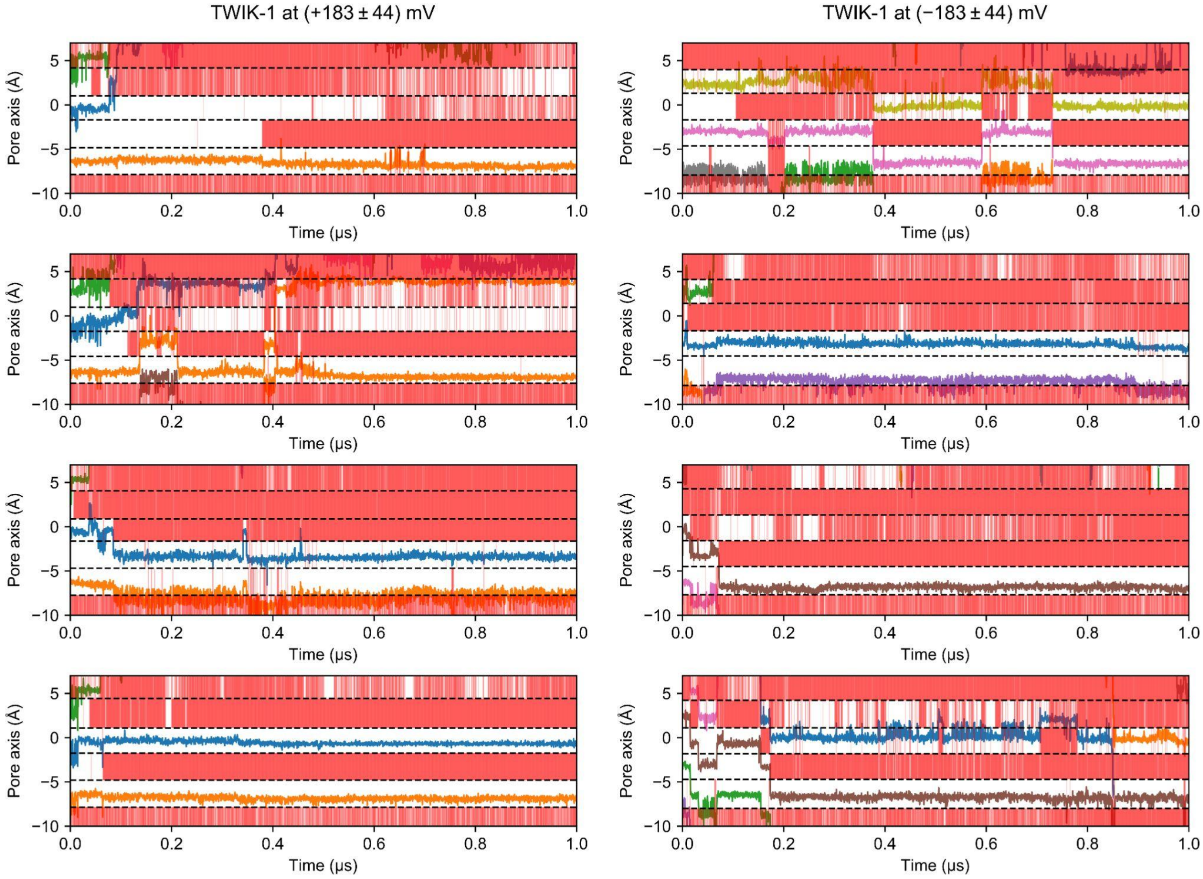


**Figure S3. Trajectories of $K^+$ ions along the pore axis in TWIK-1.** Dashed lines indicate the center of mass of the oxygen atoms delimiting the binding sites from the initial structure. Red bands within the binding sites indicate the occupancy of at least one water molecule. The voltage values are shown as mean ± SEM calculated from five 1 μs simulation runs at 303.15 K using CHARMM36m force field. Results from one simulation are shown in Fig. 2G.

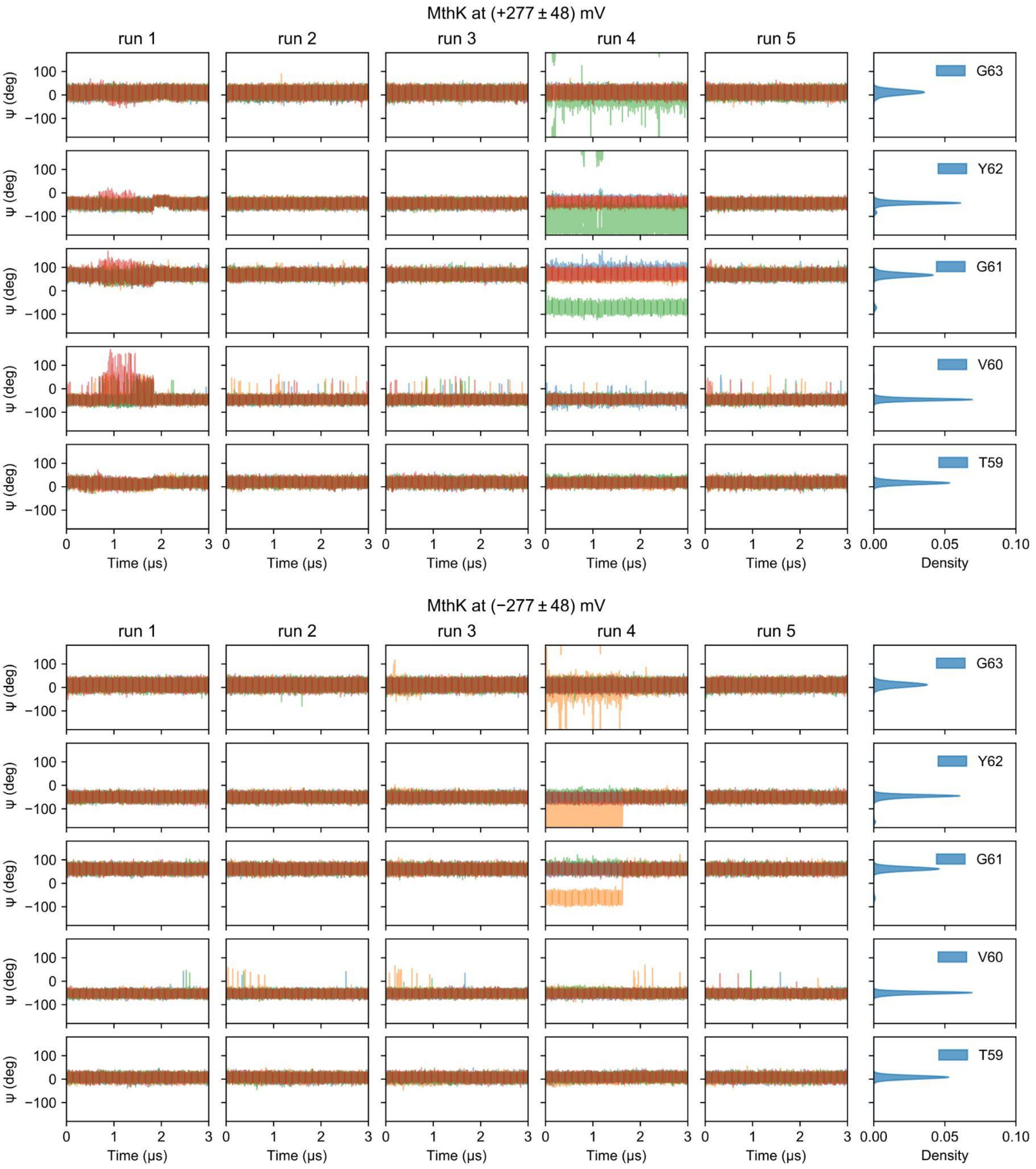


**Figure S4. Dynamics of the SF in MthK.** The ψ angle of the SF residues over time and their Gaussian kernel density estimation. Different colors indicate the residues from four pore loops. The voltage values are shown as mean ± SEM calculated from five 3 μs simulation runs at 303.15 K using CHARMM36m force field.

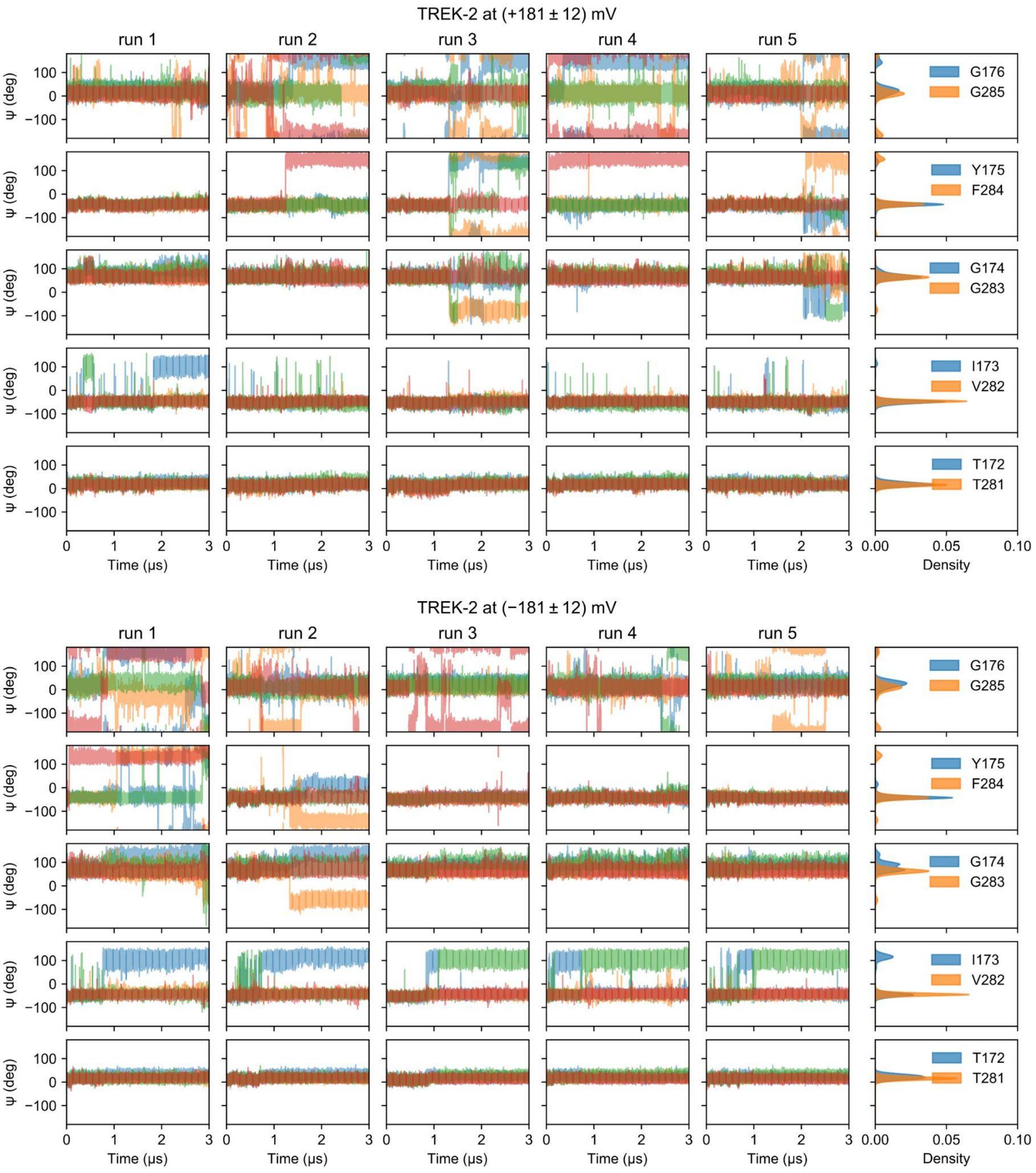


**Figure S5. Dynamics of the SF in TREK-2.** The ψ angle of the SF residues over time and their Gaussian kernel density estimation. Different colors indicate the residues from four pore loops. The voltage values are shown as mean ± SEM calculated from five 3 μs simulation runs at 303.15 K using CHARMM36m force field.

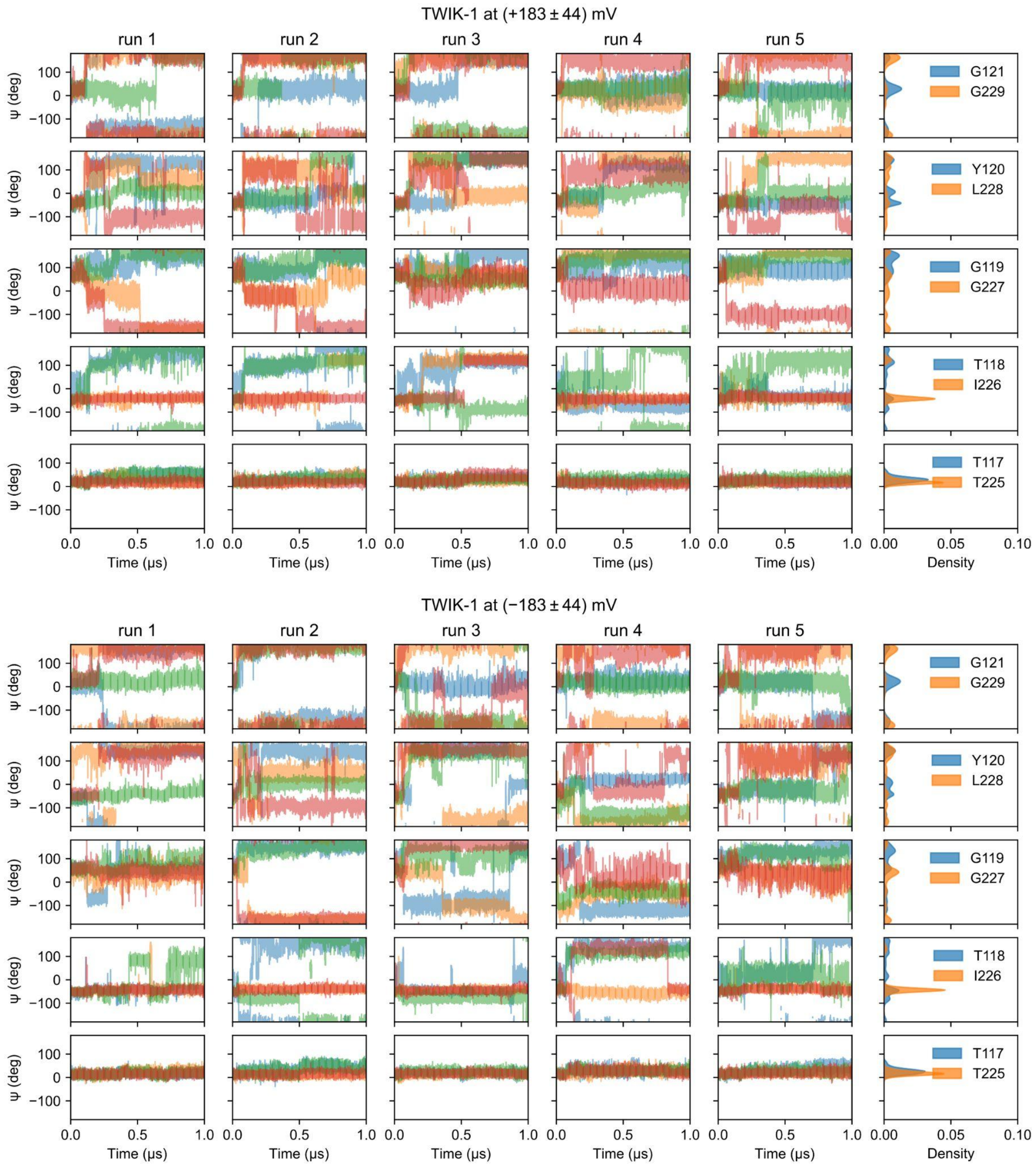


**Figure S6. Dynamics of the SF in TWIK-1.** The ψ angle of the SF residues over time and their Gaussian kernel density estimation. Different colors indicate the residues from four pore loops. The voltage values are shown as mean ± SEM calculated from five 1 μs simulation runs at 303.15 K using CHARMM36m force field.

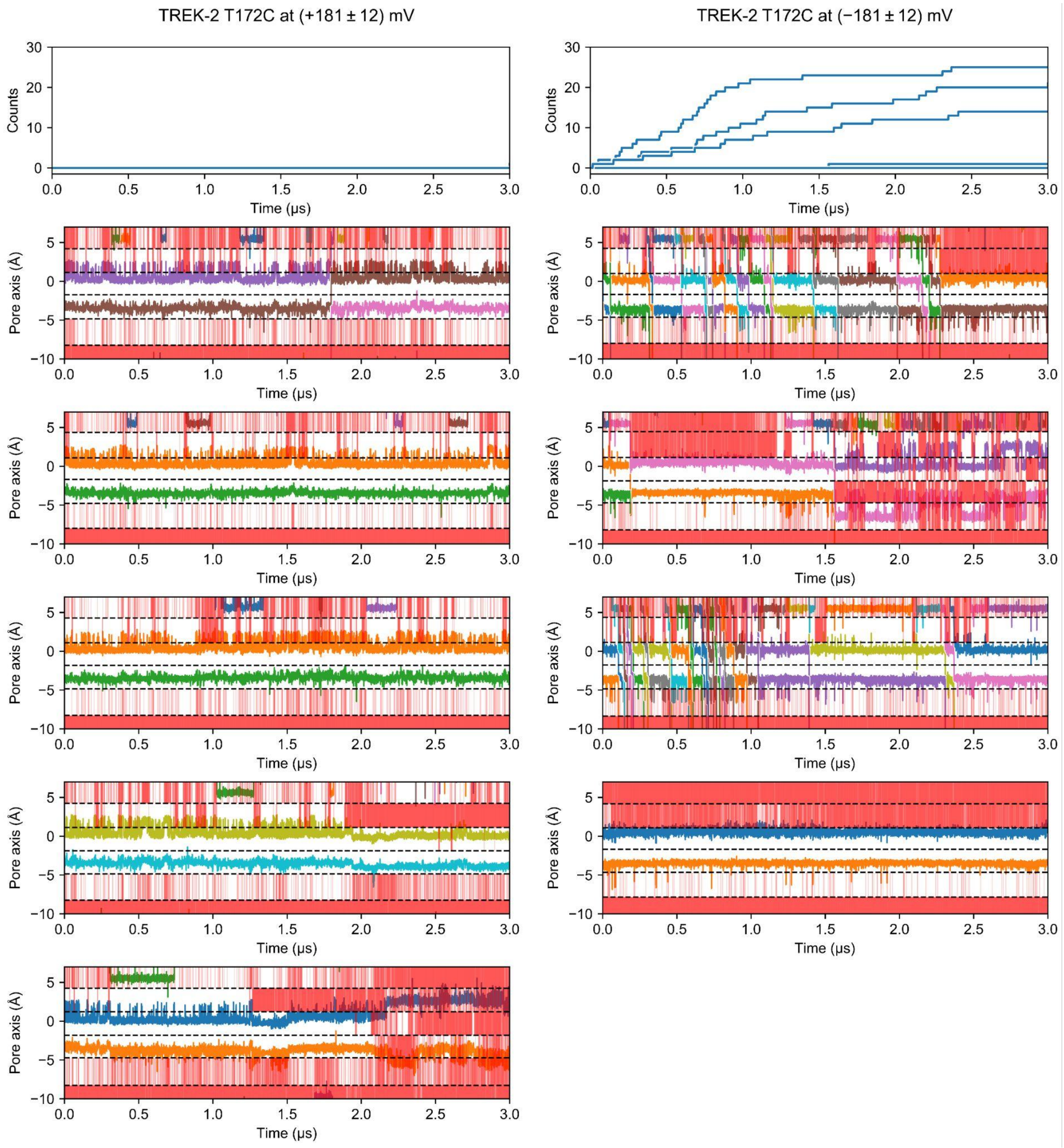


**Figure S7. Cumulative $K^+$ permeation events and trajectories of $K^+$ ions along the pore axis in TREK-2 T172C mutant.** Dashed lines indicate the center of mass of the oxygen atoms delimiting the binding sites from the initial structure. Red bands within the binding sites indicate the occupancy of at least one water molecule. The voltage values are shown as mean ± SEM calculated from five 3 μs simulation runs at 303.15 K using CHARMM36m force field. Results from one simulation under negative voltages are shown in Fig. 4A.

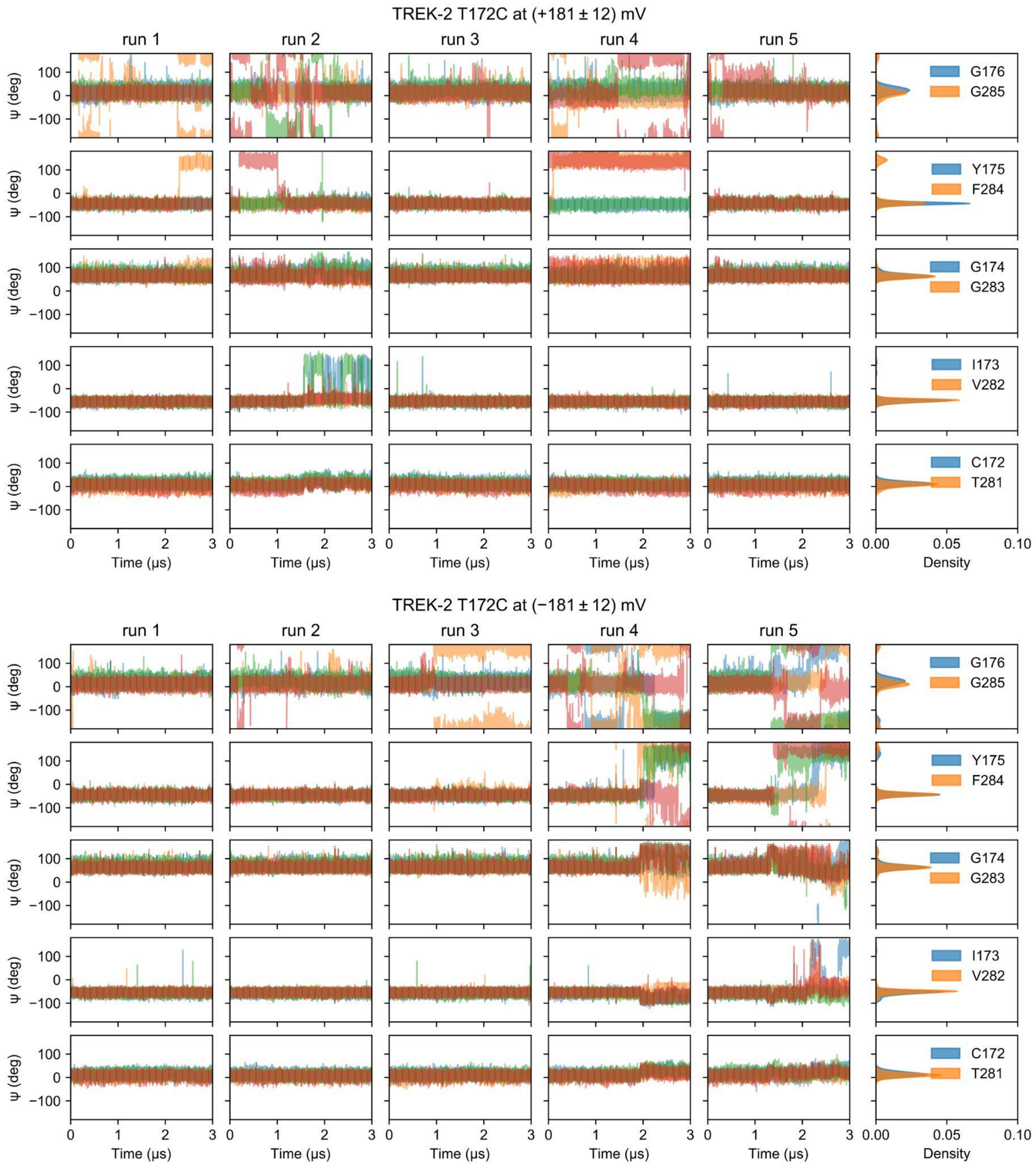


**Figure S8. Dynamics of the SF in TREK-2 T172C mutant.** The ψ angle of the SF residues over time and their Gaussian kernel density estimation. Different colors indicate the residues from four pore loops. The voltage values are shown as mean ± SEM calculated from five 3 μs simulation runs at 303.15 K using CHARMM36m force field.

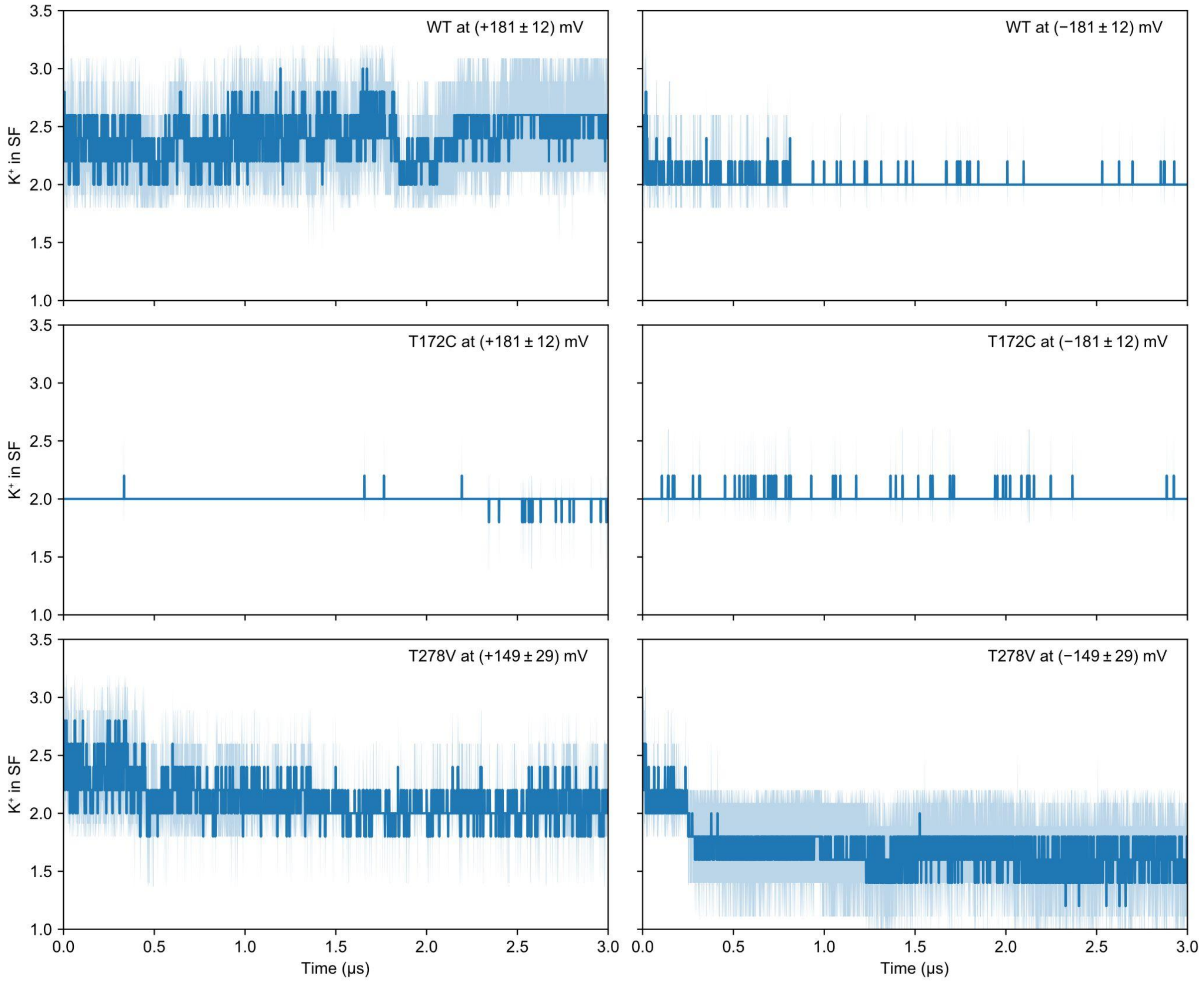


**Figure S9. Time evolution of total number of $K^+$ ions in the SF of TREK-2 WT, T172C and T278V mutants.** The voltage values are shown as mean ± SEM calculated from five 3 μs simulation runs at 303.15 K using CHARMM36m force field.

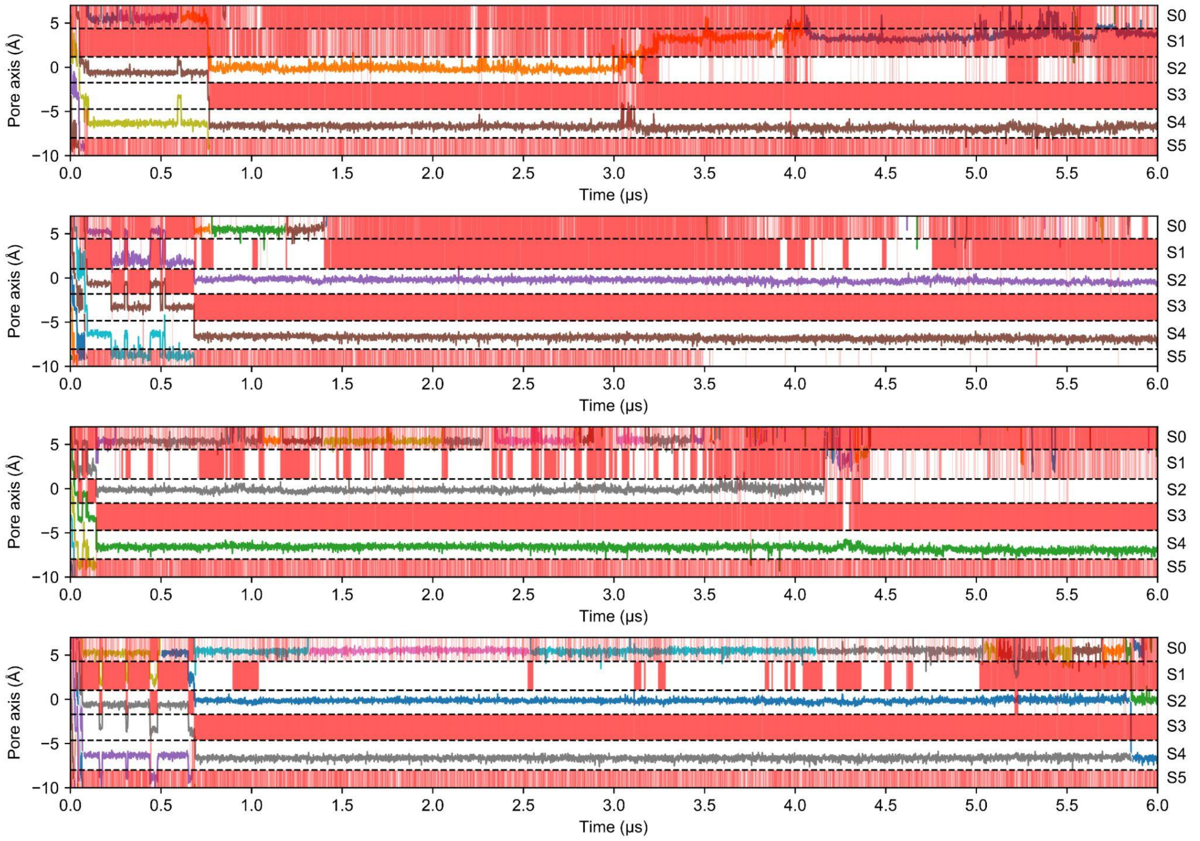


**Figure S10. Trajectories of $K^+$ ions along the pore axis in TREK-2 showing complete SF inactivation under negative voltages.** Dashed lines indicate the center of mass of the oxygen atoms delimiting the binding sites from the initial structure. Red bands within the binding sites indicate the occupancy of at least one water molecule. Simulations used the CHARMM36m force field and the system temperature was increased from 303.15 K to 323.15 K at 3 µs. Results from one simulation are shown in Fig. 5A.

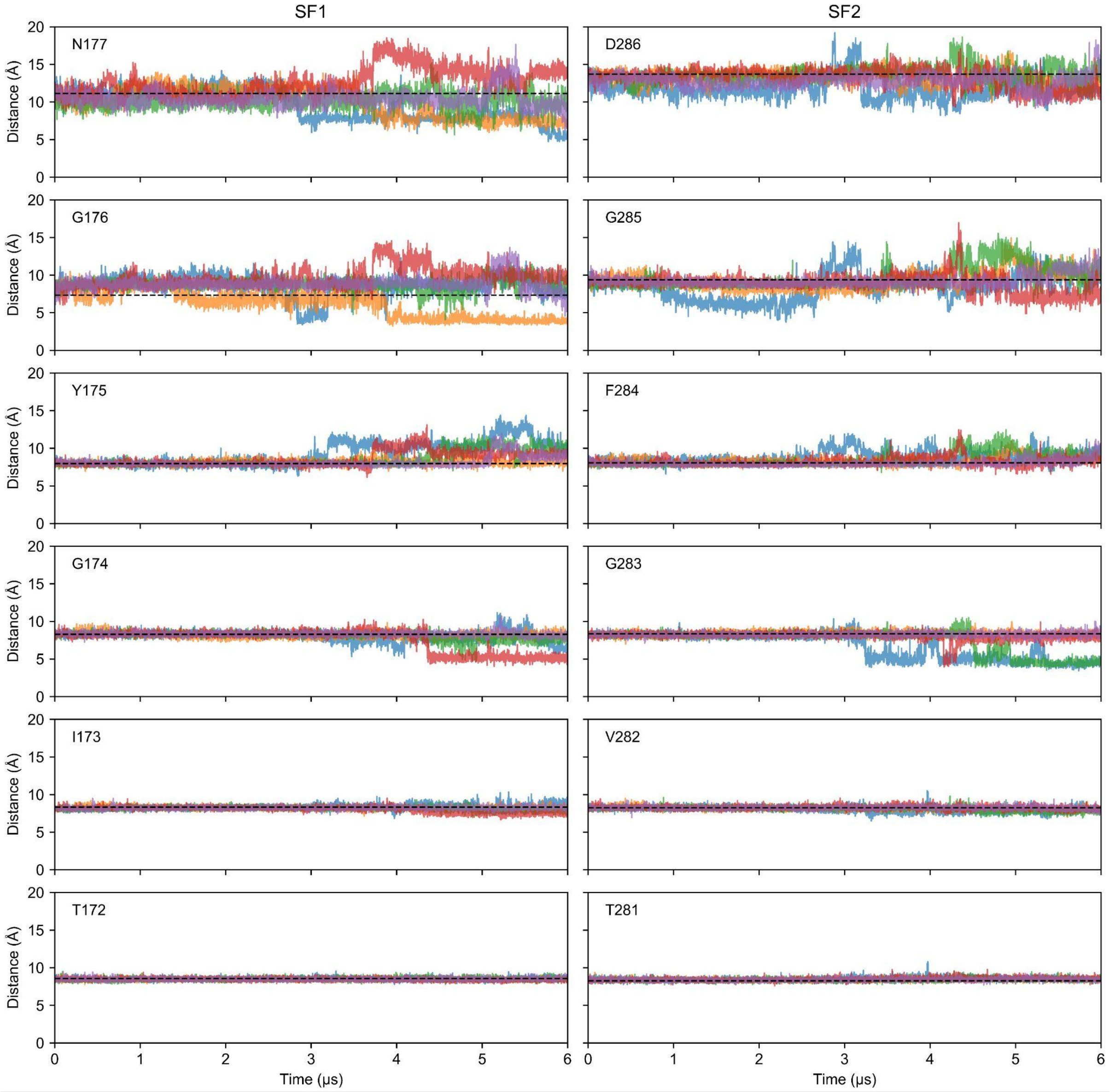


**Figure S11. Cα-Cα distances between opposing SF residues of TREK-2 during SF inactivation.** Dashed lines indicate the Cα-Cα distances from the initial structure. Different colors correspond to five simulation runs.

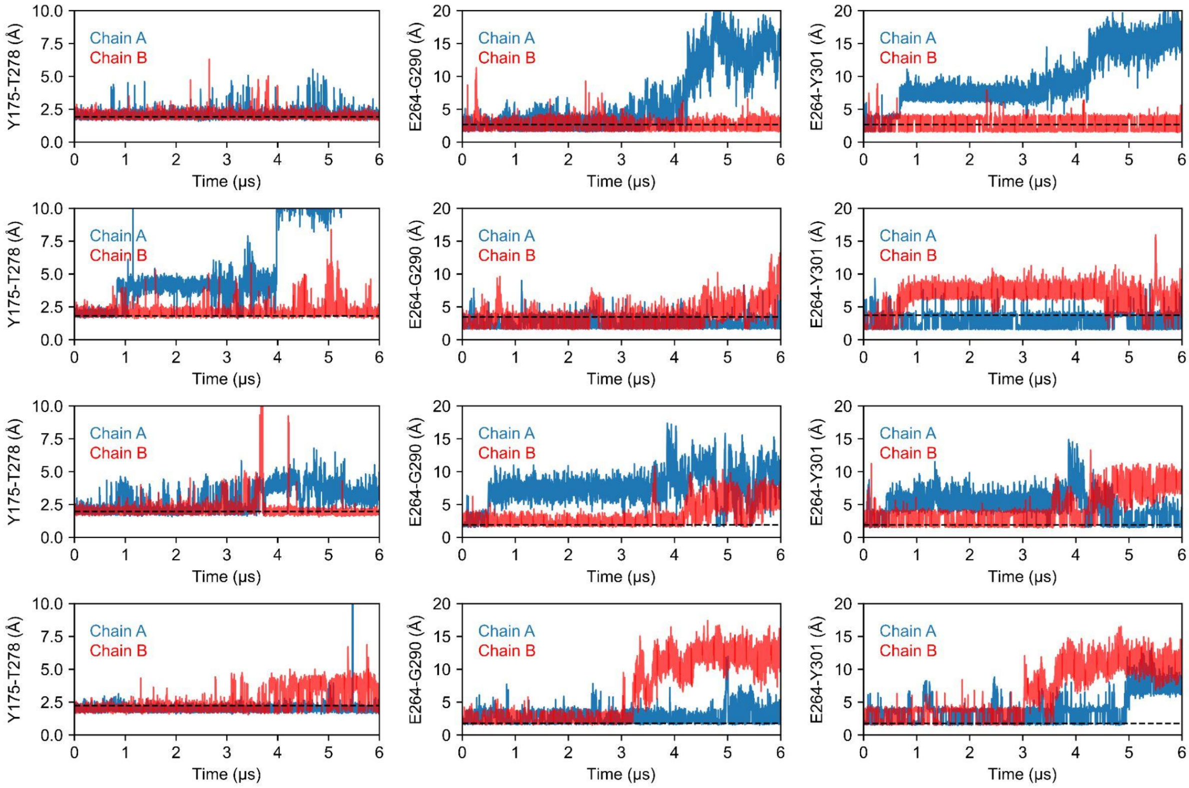


**Figure S12. Three H-bonds in TREK-2.** Distances between donor and acceptor atoms over time showing the disruption of three H-bonds during SF inactivation. Dashed lines indicate the H-bond lengths from the active state.

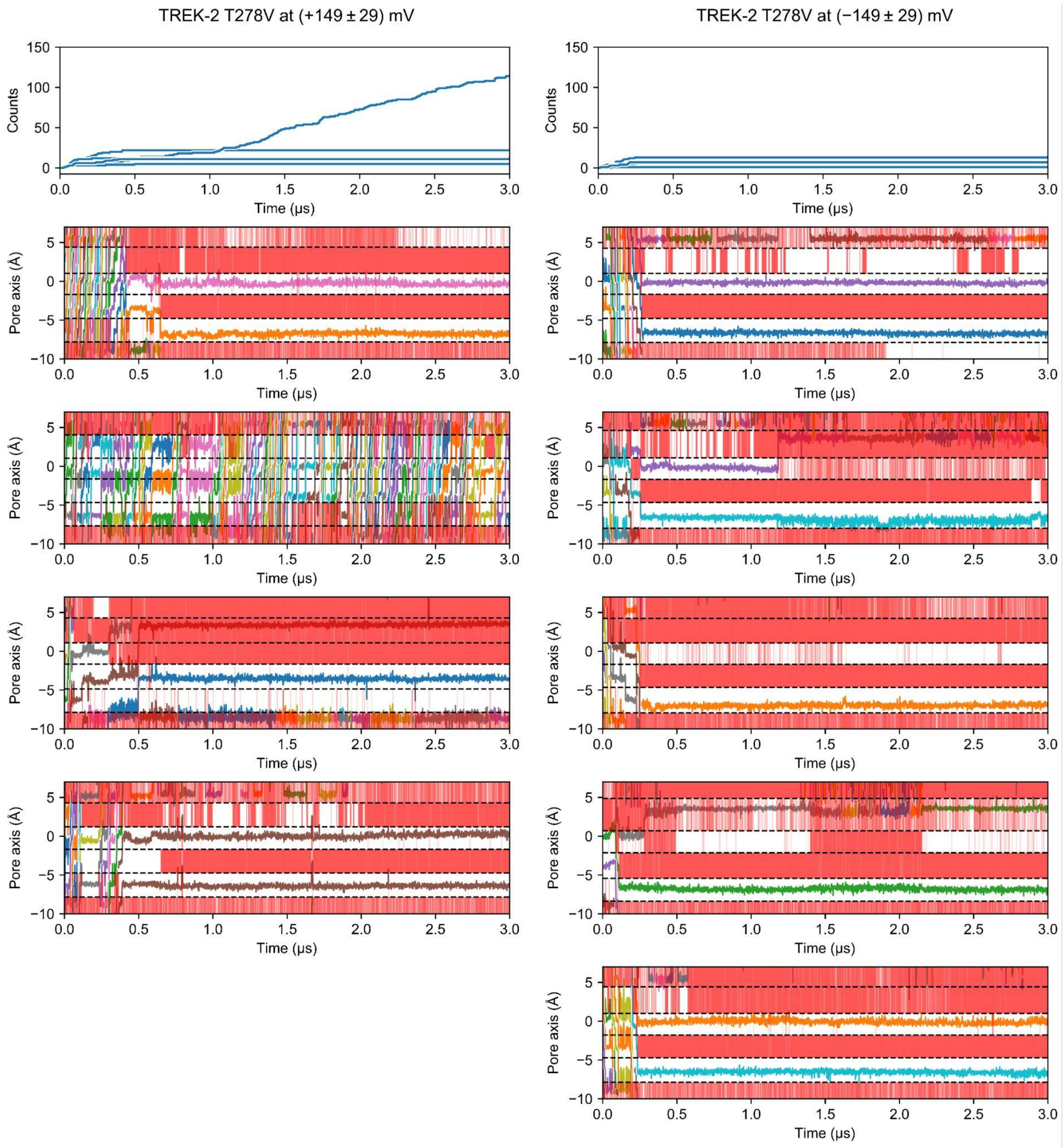


**Figure S13. Cumulative $K^+$ permeation events and trajectories of $K^+$ ions along the pore axis in TREK-2 T278V mutant.** Dashed lines indicate the center of mass of the oxygen atoms delimiting the binding sites from the initial structure. Red bands within the binding sites indicate the occupancy of at least one water molecule. The voltage values are shown as mean ± SEM calculated from five 3 μs simulation runs at 303.15 K using CHARMM36m force field. Results from one simulation under positive voltages are shown in Fig. 4D.

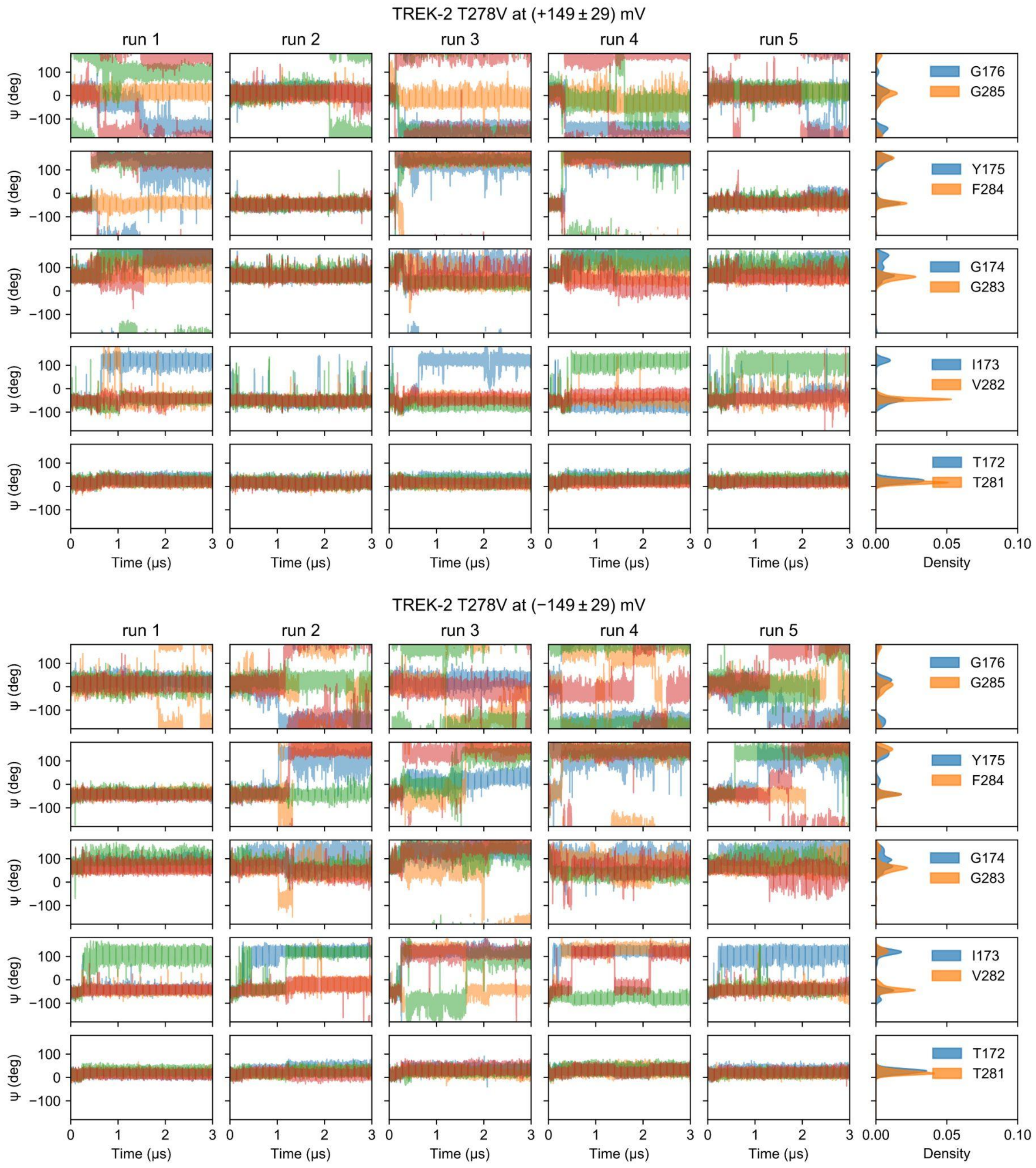


**Figure S14. Dynamics of the SF in TREK-2 T278V mutant.** The ψ angle of the SF residues over time and their Gaussian kernel density estimation. Different colors indicate the residues from four pore loops. The voltage values are shown as mean ± SEM calculated from five 3 μs simulation runs at 303.15 K using CHARMM36m force field.

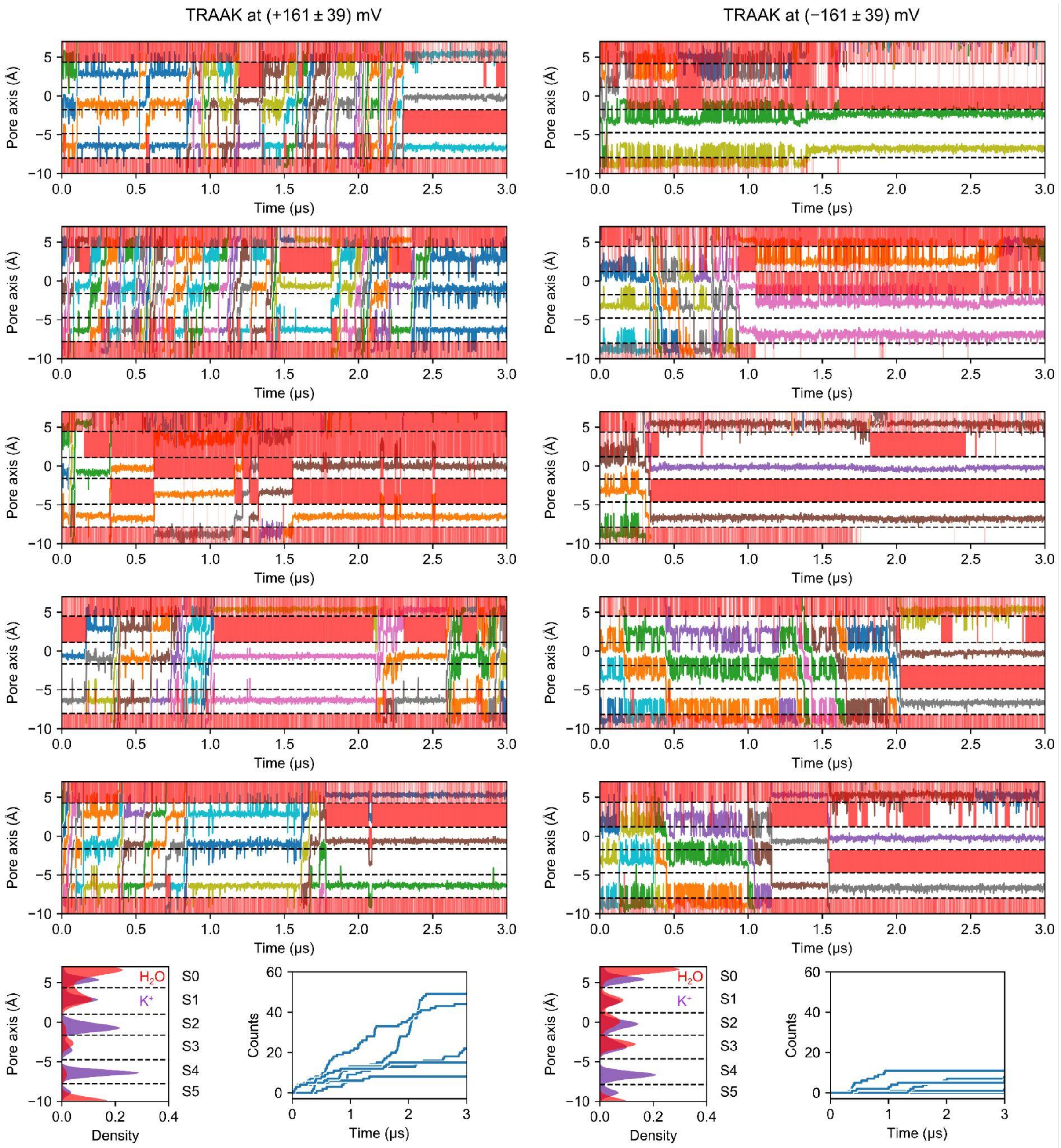


**Figure S15. Voltage-gating of TRAAK.** Trajectories of $K^+$ ions (different colors) along the pore axis in TRAAK. Dashed lines indicate the center of mass of the oxygen atoms delimiting the binding sites from the initial structure. Red bands within the binding sites indicate the occupancy of at least one water molecule. $K^+$ (purple) and water (red) density in the SF of TRAAK and cumulative $K^+$ permeation events under positive and negative voltages. The voltage values are shown as mean ± SEM calculated from five 3 μs simulation runs at 303.15 K using CHARMM36m force field.

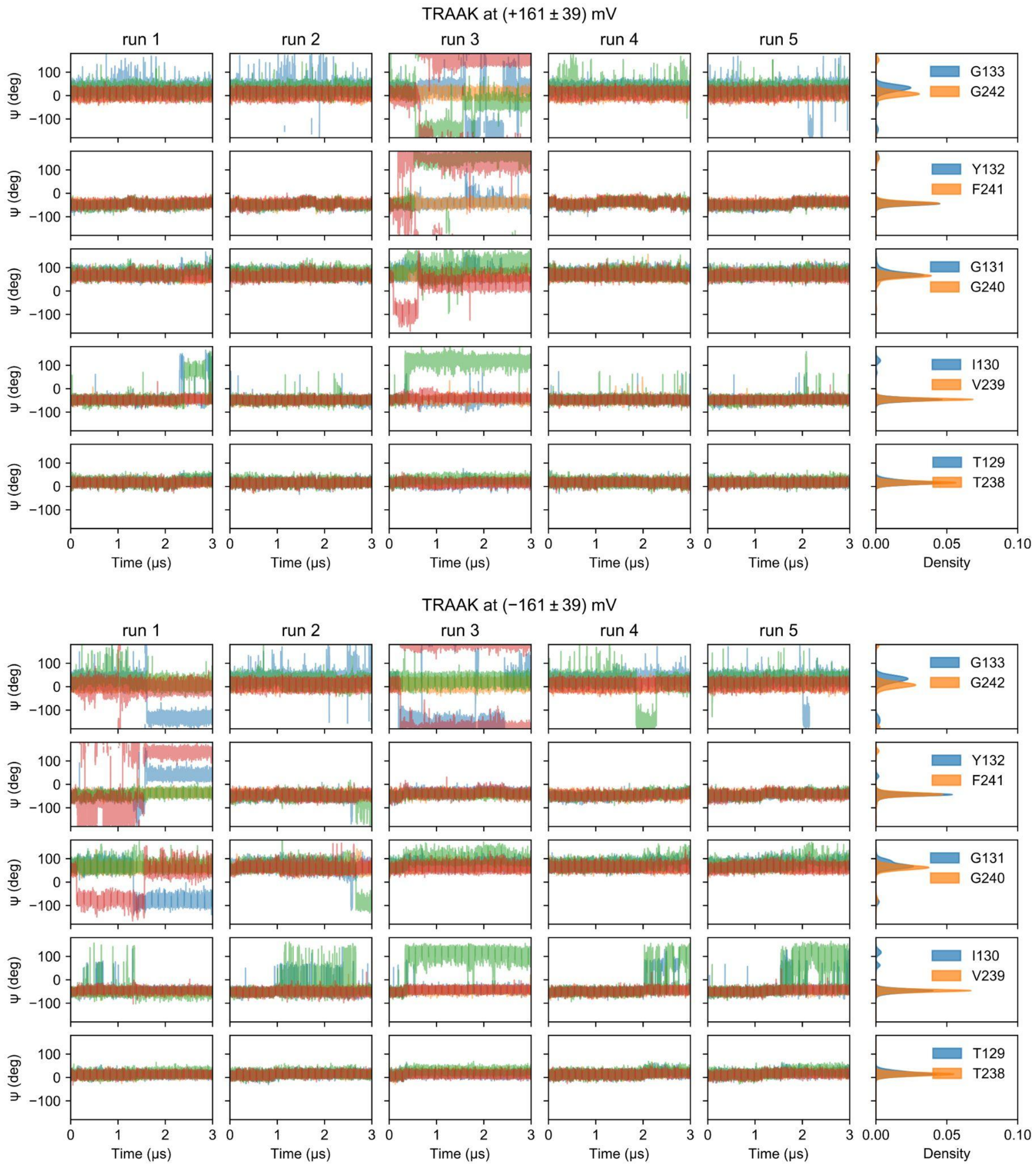


**Figure S16. Dynamics of the SF in TRAAK.** The ψ angle of the SF residues over time and their Gaussian kernel density estimation. Different colors indicate the residues from four pore loops. The voltage values are shown as mean ± SEM calculated from five 3 μs simulation runs at 303.15 K using CHARMM36m force field.

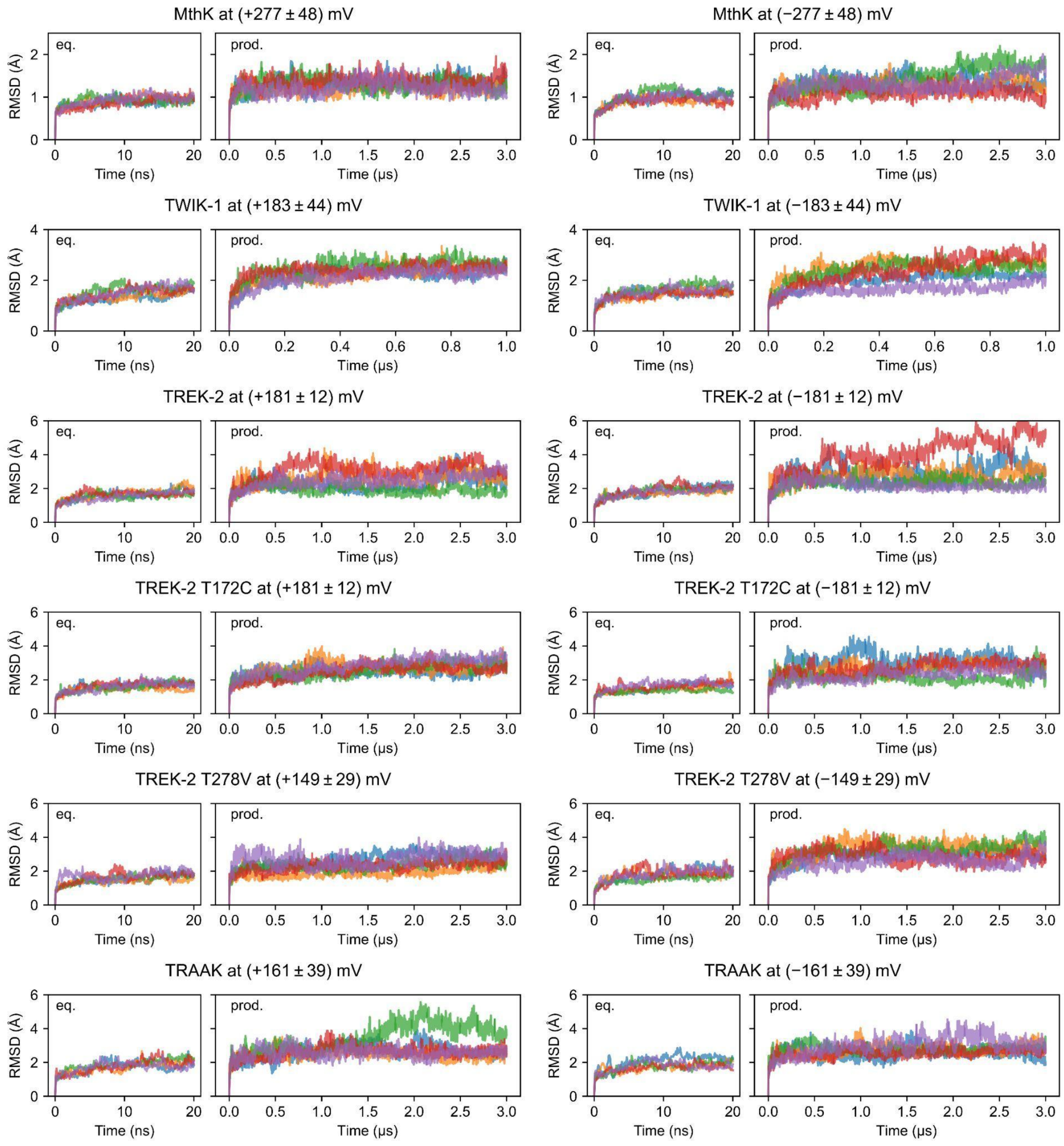


**Figure S17. RMSD of channels during equilibration and production runs.** Different colors correspond to five simulation runs. The voltage values are shown as mean ± SEM calculated from five production runs.

**Table S1.** Molecular dynamics simulations.

| System | PDB ID | N of atoms | KCl (mM) | Time (μs) | N of runs | Voltage (mV) | N of $K^+$ permeations in each run |
|---|---|---|---|---|---|---|---|
| MthK | 3LDC | 84832 | 600 | 3 | 5 | +277 ± 48 | 46, 23, 43, 0, 26 |
| | | | | | | −277 ± 48 | 9, 11, 14, 2, 6 |
| TREK-2 | 4BW5 | 235866 | 600 | 6 | 5 | +181 ± 12 | 43, 95, 63, 10, 66 |
| | | | | | | −181 ± 12 | 4, 5, 16, 3, 8 |
| TREK-2 T172C | – | 226532 | 600 | 3 | 5 | +181 ± 12 | 1, 0, 0, 0, 0 |
| | | | | | | −181 ± 12 | 21, 1, 25, 0, 14 |
| TREK-2 T278V | – | 214424 | 600 | 3 | 5 | +149 ± 29 | 22, 114, 5, 13, 12 |
| | | | | | | −149 ± 29 | 14, 4, 6, 1, 7 |
| TWIK-1 | 7SK0 | 202522 | 600 | 1 | 5 | +183 ± 44 | 0, 1, 2, 0, 0 |
| | | | | | | −183 ± 44 | 1, 1, 0, 0, 3 |
| TRAAK | 4I9W | 216904 | 600 | 3 | 5 | +161 ± 39 | 49, 44, 8, 22, 15 |
| | | | | | | −161 ± 39 | 0, 11, 2, 8, 6 |

**Movie 1.** SF1 of TREK-2 under positive voltages. The movie shows 3 μs of simulation time.

**Movie 2.** SF2 of TREK-2 under positive voltages. The movie shows 3 μs of simulation time.

**Movie 3.** SF1 of TREK-2 under negative voltages. The movie shows 3 μs of simulation time.

**Movie 4.** SF2 of TREK-2 under negative voltages. The movie shows 3 μs of simulation time.

**Movie 5.** SF1 of TREK-2 under negative voltages from the extended simulation at 323.15 K. The movie shows the last 3 μs of simulation time.

**Movie 6.** SF2 of TREK-2 under negative voltages from the extended simulation at 323.15 K. The movie shows the last 3 μs of simulation time.
